# Multivalent ubiquitin recognition regulates multiple stages of Tau self-assembly

**DOI:** 10.64898/2026.10.09.757857

**Authors:** Kevin Muwonge, Luis Fernando Durán-Armenta, Tamas Lazar, Attila Mészáros, Alexander Volkov, Álvaro M. Navarro, Cristina Marino-Buslje, Laura M. Fernández, Gustavo J. Gutierrez, Nicholas M. Kanaan, Remy Loris, Peter Tompa

**Author notes:** These authors contributed equally to this work (Kevin Muwonge, Luis Fernando Durán-Armenta). Mechanistic Biology & Profiling, The Discovery Centre, AstraZeneca, 1 Francis Crick Ave, Cambridge CB2 0AA, UK.

## Abstract

Liquid-liquid phase separation (LLPS) has emerged as an important intermediate in the pathological self-assembly of Tau, yet the molecular mechanisms that regulate this process remain incompletely understood. Although ubiquitin is a prominent component of pathological inclusions in tauopathies and other neurodegenerative diseases, its presence has largely been linked to targeting aberrant proteins for proteasomal or autophagic clearance. Here, we identify Tau as a ubiquitin-binding protein and demonstrate that multivalent, non-covalent ubiquitin recognition regulates multiple stages of Tau self-assembly. Biochemical, biophysical, structural, and computational analyses identified four ubiquitin-binding elements within the Tau microtubule-binding repeat region that engage ubiquitin’s canonical hydrophobic patch, providing a structural basis for multivalent recognition of mono- and poly-ubiquitin species. Functionally, mono- and poly-ubiquitin suppress Tau condensate formation while enhancing the liquid-like properties of condensates that persist. In contrast, mono-ubiquitin selectively enhances aggregation of the disease-linked Tau-P301L variant under conditions without detectable LLPS, demonstrating that ubiquitin exerts distinct effects depending on the molecular context and stage of Tau self-assembly. Together, these findings establish non-covalent ubiquitin recognition as a previously unrecognized mechanism regulating Tau phase behavior and aggregation. More broadly, they expand the role of ubiquitin beyond protein quality control, with ubiquitin architecture providing an additional layer of regulation over Tau condensate formation, dynamics, maturation, and aggregation.

## Introduction

Microtubule-associated protein (MAP) Tau is an intrinsically disordered protein (IDP), encoded by the *MAPT* gene and expressed as six major isoforms in the central nervous system generated through alternative splicing. These isoforms differ in the number of N-terminal inserts and microtubule-binding repeats (MTBRs). The longest isoform, Tau441 (also called 2N4R or hT40), contains four MTBRs that are essential for microtubule binding and stabilization but also contribute to Tau self-assembly and aggregation (1, 2). Tau aggregation is a hallmark of a group of neurodegenerative disorders collectively termed tauopathies, including Alzheimer’s disease (AD) and frontotemporal dementia (FTD) (3, 4). In disease, Tau undergoes conformational transitions that culminate in the formation and accumulation of insoluble amyloid assemblies, including neurofibrillary tangles, neuropil threads, and other disease-specific neuronal and glial Tau inclusions that correlate closely with neuronal dysfunction and disease progression (5–7). Although the molecular mechanisms driving the transition from soluble Tau to pathological aggregates remain incompletely understood, growing evidence suggests that liquid-liquid phase separation (LLPS) may represent an important intermediate step in Tau self-assembly (8, 9).

LLPS drives the formation of dynamic, membraneless biomolecular condensates through transient, multivalent interactions among proteins, nucleic acids, and other cellular components (10). Owing to its intrinsically disordered sequence enriched in charged and hydrophilic residues, Tau readily undergoes phase separation both *in vitro* and in cells (11, 12). Tau condensates contribute to microtubule organization and intracellular trafficking by locally concentrating tubulin and modulating motor protein interactions with the cytoskeleton (13, 14). However, disease-linked mutations and aberrant post-translational modifications (PTMs) can alter condensate material properties and promote their maturation into irreversible protein aggregates (9, 15). Recent evidence supports a mechanistic link between dysregulated phase separation and pathological aggregation, suggesting that aberrant Tau phase behavior represents a critical early event in tauopathy (8, 16).

Increasing evidence indicates that aberrant PTMs also contribute to tauopathy by altering the intermolecular interactions required for condensate formation and maturation (17–19). Notably, abnormal phosphorylation of Tau, a pathological hallmark of tauopathies, promotes phase separation, reduced microtubule affinity, and aggregation into neurofibrillary tangles (15). Other PTMs, including acetylation, are similarly implicated in Tau pathology through their effects on phase behavior, microtubule binding, and protein turnover (20–23). Ubiquitination is another PTM strongly associated with Tau dysfunction. As a covalent signal for proteasomal and autophagic degradation, ubiquitination plays a central role in Tau proteostasis. Altered Tau ubiquitination is linked to impaired proteostasis, disruption of microtubule assembly, and enhanced aggregation (24–27). Consequently, ubiquitin is a prominent component of pathological Tau inclusions in neurodegenerative disorders (28–30).

Consistent with its direct role in regulating Tau self-assembly, site-specific ubiquitination within the MTBRs modulates condensate formation in a site- and cofactor-dependent manner (31). Beyond its canonical role as a covalent protein modifier, ubiquitin recently emerged as a regulator of biomolecular condensates through reversible, non-covalent interactions.

Ubiquitin-mediated interactions modulate the phase behavior of several proteins associated with neurodegenerative diseases, including ubiquilin-2 (UBQLN2) and sequestosome-1 (SQSTM1/p62) (32). However, whether Tau directly interacts with free ubiquitin and how these interactions influence phase separation and aggregation remain unknown.

In this study, we present the structural basis and functional consequences of non-covalent Tau–ubiquitin interactions using interactome analysis, biophysical characterization, structural modeling, molecular dynamics simulations, and *in vitro* phase separation and aggregation assays. Using wild-type Tau441 and the FTD-associated P301L variant, we identified four ubiquitin-binding elements within the microtubule-binding repeat region that mediate multivalent ubiquitin recognition. We further show that these interactions suppress condensate formation, preserve liquid-like condensate material properties, and differentially regulate amyloid aggregation. Together, our findings identify ubiquitin as a direct regulator of Tau phase separation and aggregation through multivalent non-covalent interactions within the MTBR.

## Results

### Interactome and biochemical analyses identify Tau as a ubiquitin-binding protein

Ubiquitinated protein inclusions are a pathological hallmark of several neurodegenerative diseases. To identify candidate ubiquitin-binding proteins associated with these inclusions, we analyzed the amyotrophic lateral sclerosis (ALS) inclusion body proteome. The 52 proteins identified within ALS-linked inclusions (33) form a densely interconnected protein-protein interaction network **(Supporting Fig. S1A)** that is significantly enriched compared with randomly sampled human protein networks of equal size generated from BioGRID and IntAct interactomes **(Fig. 1A; Supporting Fig. S1B–C)**.

**Figure 1.**
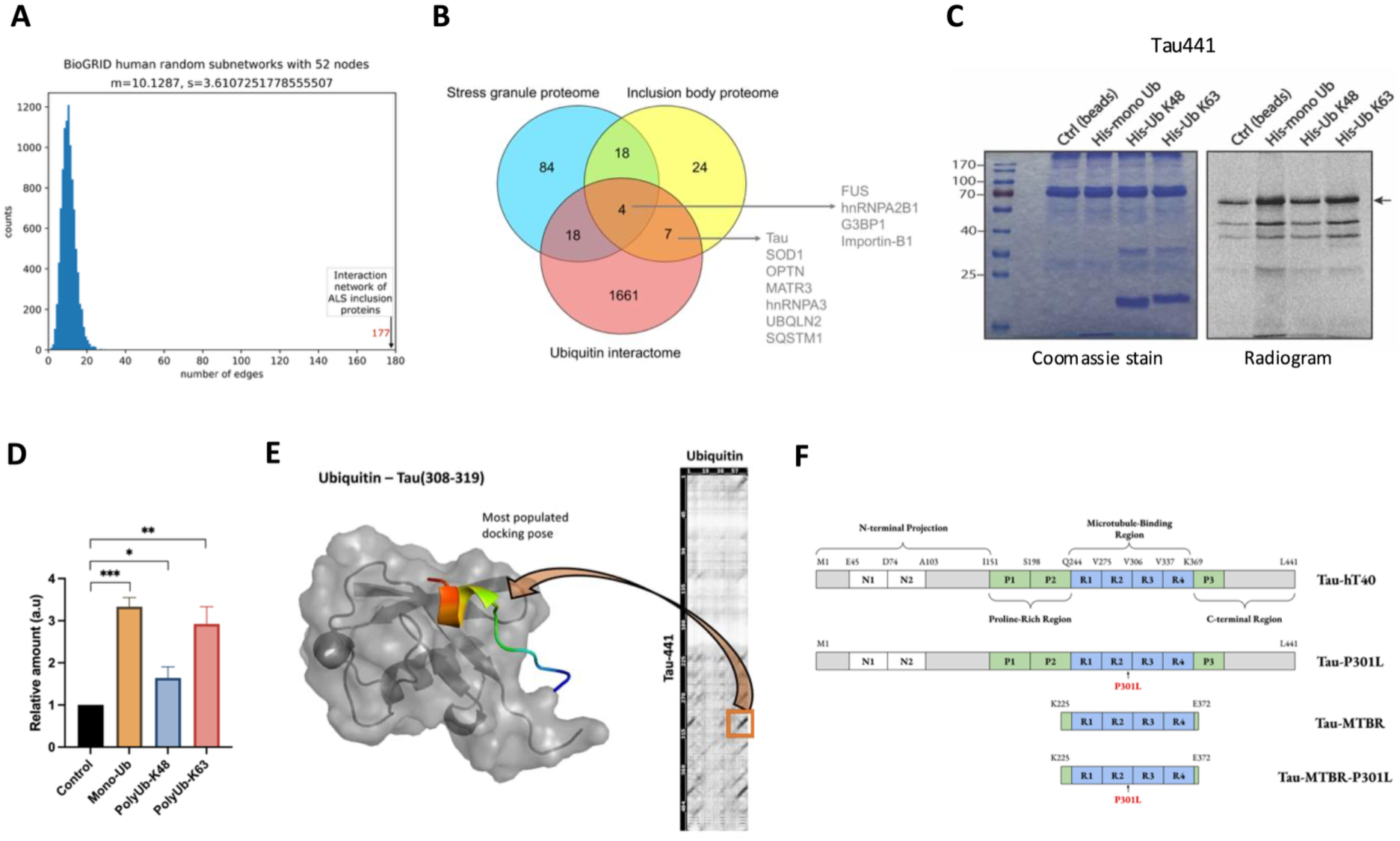
Identification and validation of Tau as a ubiquitin-binding protein. **(A)** Protein–protein interaction connectivity of the ALS inclusion body proteome compared with 10,000 randomly sampled human protein networks of the same size, demonstrating significant enrichment in network connectivity. (**B**) Integration of the ALS inclusion body proteome, the stress granule proteome, and the BIOGRID ubiquitin interactome identifies candidate ubiquitin-binding proteins, including Tau. **(C)** Experimental validation of Tau ubiquitin binding by pull-down assays using ^35^S-radiolabeled Tau and different ubiquitin probes. **(D)** Densitometric quantification of Tau recovered from three independent pull-down experiments identifies specific ubiquitin binding (graph represents mean ± SD; one-way ANOVA with Sidak’s multiple comparisons test; n = 3 independent biological replicates; \*\*\**p* = 0.0005, \*\**p* = 0.0012, \**p* = 0.0169). **(E)** Docking predicts the binding of Tau’s ubiquitin-interacting motif (residues ^308^IVYKPVDLSKVT^319^) to the canonical hydrophobic patch of ubiquitin, while co-evolution analysis independently supports the predicted interaction interface. **(F)** Domain organization of the longest human Tau isoform (Tau441; hT40) and the recombinant Tau constructs used throughout this study. The P301L mutation was introduced into both Tau441 and Tau-MTBR constructs.

The ALS inclusion proteome contains 117 interaction edges, far exceeding the 10 ± 3.6 edges expected by chance, indicating that these proteins constitute a highly interconnected functional network. Integration of the ALS inclusion body proteome with the stress granule proteome (34) and the ubiquitin interactome (35) identified eleven candidate ubiquitin-binding proteins, all of which are found in heavily ubiquitinated inclusions in post-mortem ALS/FTD patient tissue (36, 37) **(Fig. 1B)**. Among the candidates, Tau emerged as a previously unrecognized putative ubiquitin-binding protein.

To validate the computational predictions, interactions between the eleven candidates and mono-ubiquitin, Lys48 (K48)-linked di-ubiquitin, or Lys63 (K63)-linked di-ubiquitin were assessed by pull-down assays **(Supporting Fig. S2)**. Established ubiquitin-binding proteins, including SQSTM1/p62, ubiquilin-2, and optineurin, robustly bind all three ubiquitin species, confirming assay specificity. In contrast, G3BP1, FUS, and matrin-3 showed no detectable interaction, whereas hnRNPA2, hnRNPA3, and importin-β1 exhibit selective or weaker binding profiles **(Supporting Fig. S2)**. Notably, full-length Tau441 robustly bound mono-ubiquitin and K63-linked di-ubiquitin, while exhibiting weaker interaction with K48-linked di-ubiquitin **(Fig. 1C–D)**. Direct Tau–ubiquitin interaction was further confirmed by biolayer interferometry (BLI), which yields an apparent dissociation constant (*K*_D_) of 260 ± 117 µM **(Supporting Fig. S3)**. Together, these findings establish Tau as a ubiquitin-binding protein and provide the foundation for subsequent structural and functional analyses.

To identify the region of Tau responsible for ubiquitin recognition, we combined sequence analysis, structural modeling, and coevolutionary approaches **(Supporting Fig. S4)**. Charge distribution analysis reveals a highly positively charged central region encompassing the proline-rich domain and the microtubule-binding repeat region (MTBR), whereas both termini are predominantly negatively charged **(Supporting Fig. S4A)**. Molecular docking, co-evolution analysis, and mapping of known ubiquitin-binding surfaces consistently predict that Tau interacts with the canonical hydrophobic patch of ubiquitin through residues within its MTBR (^308^IVYKPVDLSKVT^319^) **(Fig. 1E; Supporting Fig. S4B)**, despite lacking a disordered ubiquitin-binding motif (DisUBM) (38).

AlphaFold-Multimer independently supports this interaction mode, placing an MTBR-derived peptide (^305^SVQIVYKPVD^314^) against the ubiquitin hydrophobic patch in the highest-confidence models **(Supporting Fig. S4C)**. A similar binding geometry is also predicted for the small ubiquitin-like modifiers, SUMO1 and SUMO2 **(Supporting Fig. S5)**. Together, these analyses converge on the MTBR as the most likely ubiquitin-binding region within Tau. To test these predictions experimentally, we generated a panel of full-length and MTBR-derived constructs, including the FTD-associated P301L variant, for biophysical, structural, phase separation, and aggregation studies **(Fig. 1F)**.

### Tau harbors four distinct ubiquitin-binding elements within its MTBR

Nuclear magnetic resonance (NMR) spectroscopy experimentally validates the predicted Tau–ubiquitin interaction interface and confirms that Tau recognizes the canonical hydrophobic patch of ubiquitin. Titration of unlabeled Tau-MTBR (K18) into ^15^N-labeled ubiquitin produces localized chemical shift perturbations (CSPs), consistent with direct binding between the two proteins **(Fig. 2A)**. Mapping the average CSPs (Δδ_avg_) onto the ubiquitin structure reveals that the most strongly perturbed residues (colored in red) cluster around the canonical hydrophobic patch centered on Ile44 and Val70 **(Fig. 2B)**. Smaller CSPs (colored in yellow) extend along the surrounding surface, consistent with dynamic interactions expected for an intrinsically disordered protein. Simultaneous fitting of CSPs from representative ubiquitin residues (K6, A46, H68 and L69) yields a *K*_D_ of 220 ± 39 µM **(Supporting Fig. S6A)**. An independent microscale thermophoresis (MST) assay produced a comparable affinity **(Supporting Fig. S6B)**, providing orthogonal validation of the interaction.

**Figure 2.**
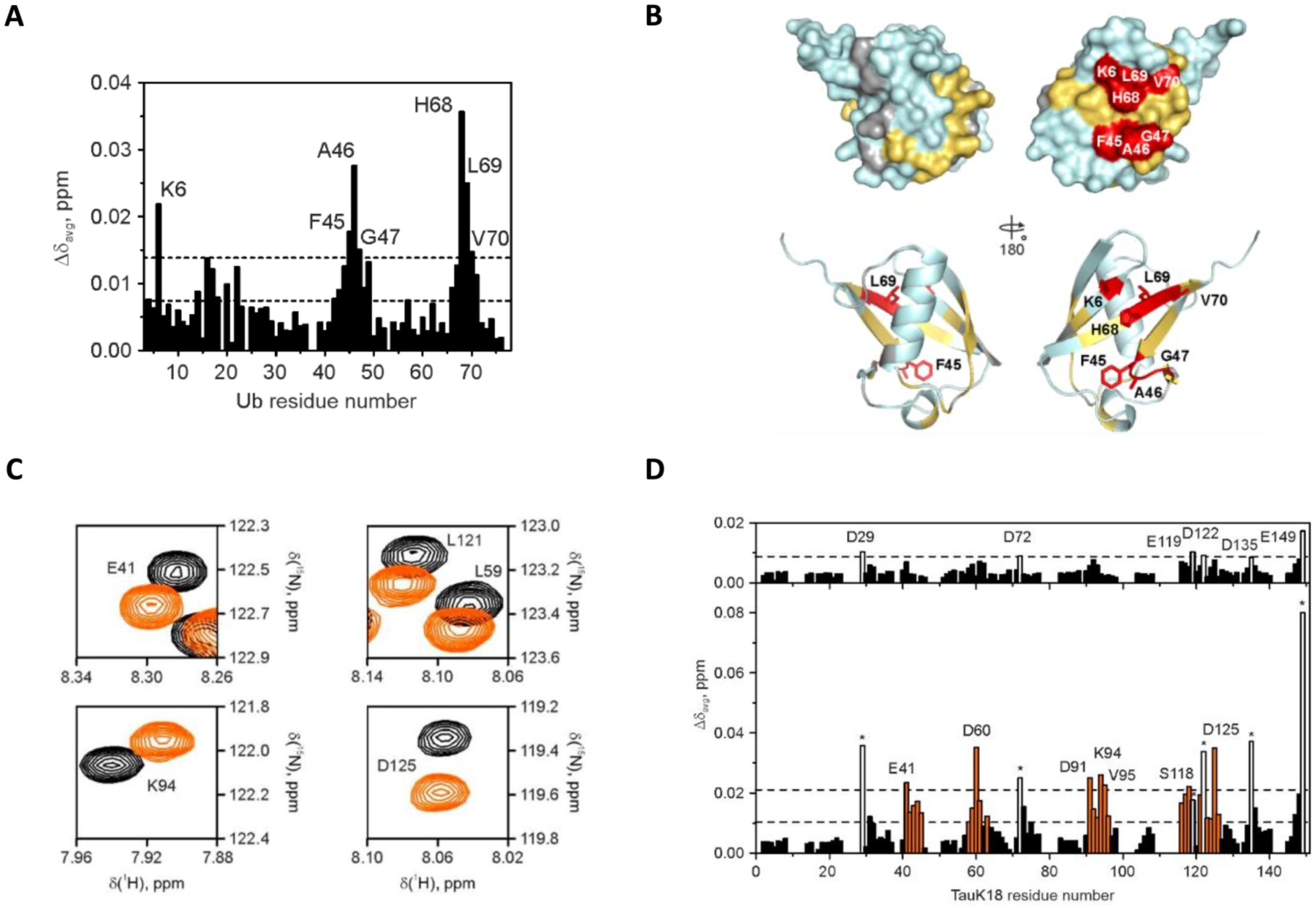
NMR mapping of the Tau–ubiquitin interaction reveals the molecular binding interface. **(A)** NMR chemical shift perturbation (CSP) analysis identifies ubiquitin residues perturbed upon binding to Tau-MTBR. Average backbone amide chemical shift changes (Δδ_avg_) of ^15^N-labeled ubiquitin were measured after adding three molar equivalents of Tau-MTBR. Horizontal dashed lines indicate the average Δδ_avg_ and the average (avg) plus one standard deviation (SD). **(B)** Mapping of the observed CSPs onto the ubiquitin structure (PDB: 1UBQ) reveals a distinct Tau-binding interface. Residues exhibiting Δδ_avg_ values greater than the mean are highlighted (red: Δδ_avg_ > avg + SD; yellow: Δδ_avg_ > avg). Prolines and residues with unassigned resonances are shown in gray, while residues exhibiting the largest Δδ_avg_ are labeled and displayed as sticks in the bottom view. (**C**) Reciprocal NMR titration of [^13^C, ^15^N]-labeled Tau-MTBR with three molar equivalents of ubiquitin identifies Tau residues involved in ubiquitin recognition. Representative Tau-MTBR backbone amide resonances (Δδ_avg_) are shown for free Tau-MTBR (black) and the Tau-MTBR–ubiquitin complex (orange). **(D)** Average backbone amide CSPs (Δδ_avg_) across Tau-MTBR identify contiguous ubiquitin-binding regions spanning four or more residues, highlighted in orange. Residues experiencing the largest effects (Δδ_avg_ > avg + 2 SD) are indicated by open bars and asterisks in the bottom panel and labeled in the top panel.

Reciprocal titrations using [^13^C,^15^N]-labeled Tau-MTBR and unlabeled ubiquitin identify four distinct ubiquitin-binding elements distributed across the four pseudo-repeats (R1–R4) of Tau’s MTBR **(Fig. 2C–D)**. The R1 element (^264^ENLKH^268^) represents a previously unrecognized ubiquitin-binding site, whereas the R2 element (^281^KLDLSNV^287^) lies adjacent to the PHF6* aggregation motif (^275^VQIINK^280^). The strongest perturbations are observed within the R3 element (^314^DLSKV^318^), which lies adjacent to the PHF6 aggregation motif (^306^VQIVYK^311^), separated by residues ^312^PV^313^, and coincides with the interaction interface independently predicted by molecular docking, coevolution analysis, and AlphaFold-Multimer **(Fig. 1E; Supporting Fig. S4B-C).** A fourth binding element (^339^VKSEKLDFKDR^349^) is identified within R4 and is located adjacent to the recently described Polymorphic Amyloid Motif of Repeat 4 (PAM4; ^350^VQSKIGSLDNITH^362^) that is implicated in Tau amyloid assembly (39). Interestingly, three of the four ubiquitin-binding elements are positioned adjacent to established amyloidogenic regions within the MTBR.

Together, these findings demonstrate that Tau recognizes ubiquitin through four spatially separated binding elements within the MTBR, establishing a structural basis for multivalent recognition of mono- and poly-ubiquitin species. The presence of multiple binding sites is consistent with the stronger interaction observed with mono-ubiquitin and K63-linked ubiquitin species, which predominantly adopt extended conformations with accessible surfaces (40), relative to K48-linked ubiquitin chains, which result in more compact conformations **(Fig. 1C).**

### Molecular dynamics simulations reveal multivalent Tau recognition of poly-ubiquitin

Having identified four ubiquitin-binding elements within the Tau MTBR, we used molecular dynamics (MD) simulations to investigate how two sites, Patch 1 (Tau repeat R2, ^281^KLDLSN^286^) and Patch 2 (Tau repeat R3, ^314^DLSKVT^319^), could cooperate in binding head-to-tail linked di-ubiquitin. Starting from an NMR-guided docking model, we performed two independent simulations of the Tau R1–R4/di-ubiquitin complex. In replica 1, the complex dissociates at approximately 270 ns **(Supporting Movie M1)**, whereas in replica 2, it remains stably associated throughout the full trajectory **(Supporting Movie M2)**.

Tau binding elements maintained dynamic contact with the canonical hydrophobic patches of both ubiquitin moieties, with dissociation at one contact often compensated by interaction at another site, rather than reliance on a single interface **(Fig. 3A & Supporting Movie M2)**. Because we extrapolated the mono-ubiquitin NMR restraints to the second ubiquitin moiety, the resulting bivalent geometry represents a predictive model rather than an experimentally determined interaction mode. Overall, the simulations are consistent with Tau’s fuzzy binding to ubiquitin chains and support the relatively weak interaction affinity of single valencies.

**Figure 3.**
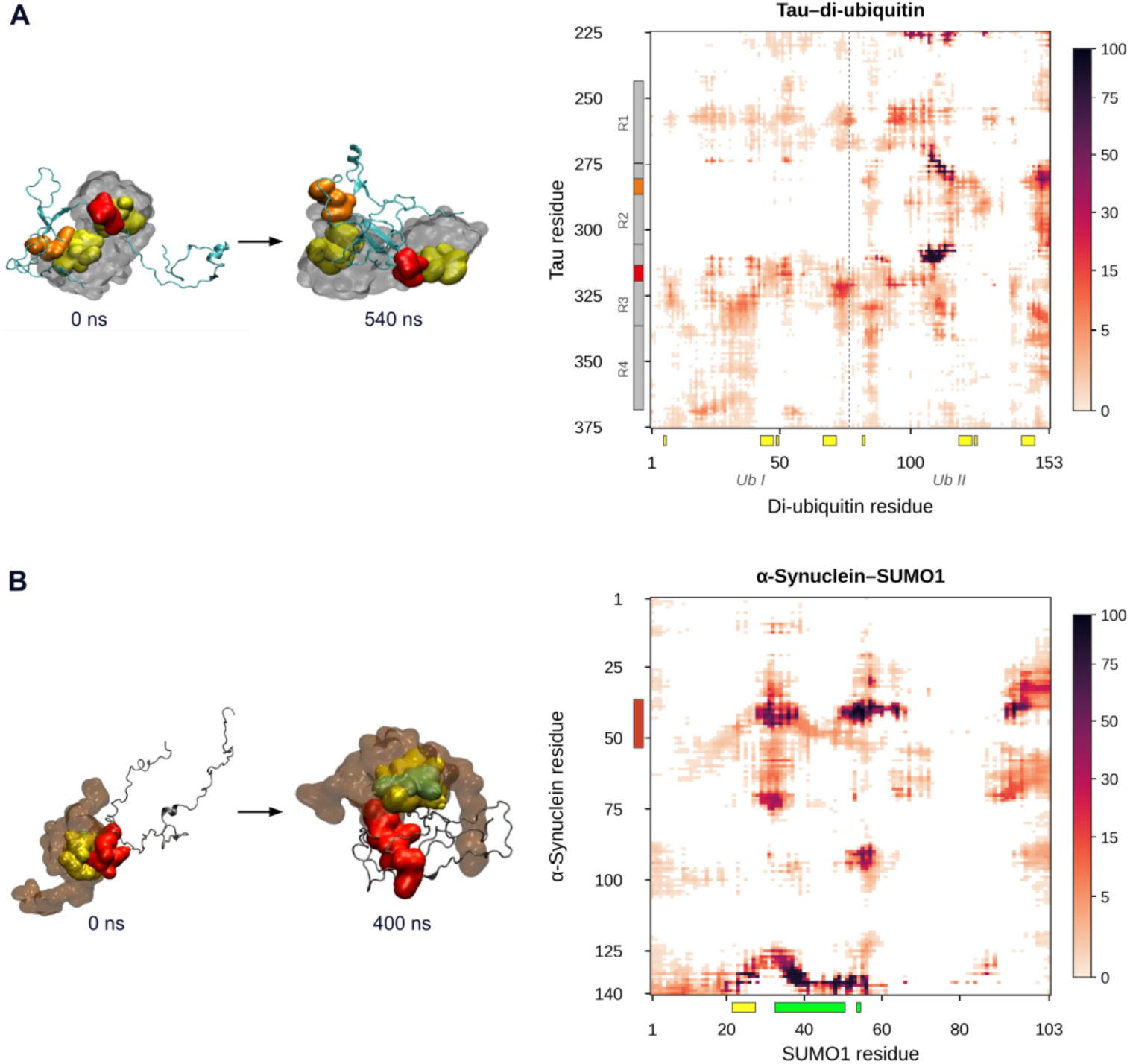
Distinct binding modes of Tau–di-ubiquitin and α-synuclein–SUMO1 revealed by molecular dynamics. (**A**) Snapshots of the Tau R1–R4/di-ubiquitin complex at 0 ns and 540 ns from the replica shown in **Supporting Movie M2**. Tau is shown in cyan and di-ubiquitin as a gray surface, with Patch 1 (P1, ^281^KLDLSN^286^) and Patch 2 (P2, ^314^DLSKVT^319^) highlighted in orange and red, respectively. NMR-defined ubiquitin interface residues (K6, ^43^LIFAG^47^, Q49, and ^67^HLVLR^71^) are shown in yellow on both ubiquitin moieties. The adjoining contact map shows Tau (Y-axis) versus di-ubiquitin (X-axis) and uses 7.4 Å as the threshold for considering two residues to be interacting. The gray bar denotes Tau repeats R1–R4, with P1/P2 patches highlighted in orange/red, and yellow boxes on the X-axis indicate the ubiquitin interface residues. The dashed line separates the Ub I (M1–G76) and Ub II (M77–G153) linked head-to-tail. **(B)**. Snapshots of the α-synuclein–SUMO1 complex at 0 ns and 400 ns from the replica shown in **Supporting Movie M3**. SUMO1 is shown as a tan surface and α-synuclein as a gray cartoon. The adjoining contact map shows α-synuclein (Y-axis) versus SUMO1 (X-axis), with docking restraints from (41) indicated for α-synuclein residues G37–V53 (red) and SUMO1 residues H22–V27 (yellow), K33–V50, and E54 (green). For both contact maps, residue pairs with a minimum distance <7.4 Å were considered in contact, and color indicates contact occupancy (percentage of simulation frames in contact), from 0 (white) to 100%.

As a reference system, we similarly simulated the reported α-synuclein–SUMO1 complex (41) in two independent replicas. Consistent with this work, SUMO1 binding was accompanied by α-synuclein compaction: the complex remained associated in replica 1 throughout the full 400 ns simulation **(Fig. 3B & Supporting Movie M3)** but dissociated at approximately 90 ns in replica 2 **(Supporting Movie M4)**. By comparison, α-synuclein also maintained multiple simultaneous contacts with SUMO1, consistent with a similarly dynamic multivalent binding mechanism; however, in the case of Tau, this was accompanied by a comparatively modest global compaction. Together, these simulations support a model in which multiple Tau binding elements dynamically engage distinct ubiquitin moieties, but do not fully match those observed for homolog SUMO1 with its IDP partner, providing a potentially new structural mechanism for IDP recognition of ubiquitin and ubiquitin-like modifiers.

### Mono- and poly-ubiquitin differentially regulate Tau liquid-liquid phase separation

To determine the functional consequences of Tau–ubiquitin interactions, we established an *in vitro* liquid-liquid phase separation (LLPS) assay at room temperature using physiological Tau concentrations (4 µM) under molecular crowding conditions (10% PEG). Green fluorescent protein (GFP)-tagged wild-type Tau (Tau-GFP) and the FTD-associated mutant Tau-P301L-GFP readily formed spherical condensates that increased in size and abundance over time **(Fig. 4A).**

**Figure 4.**
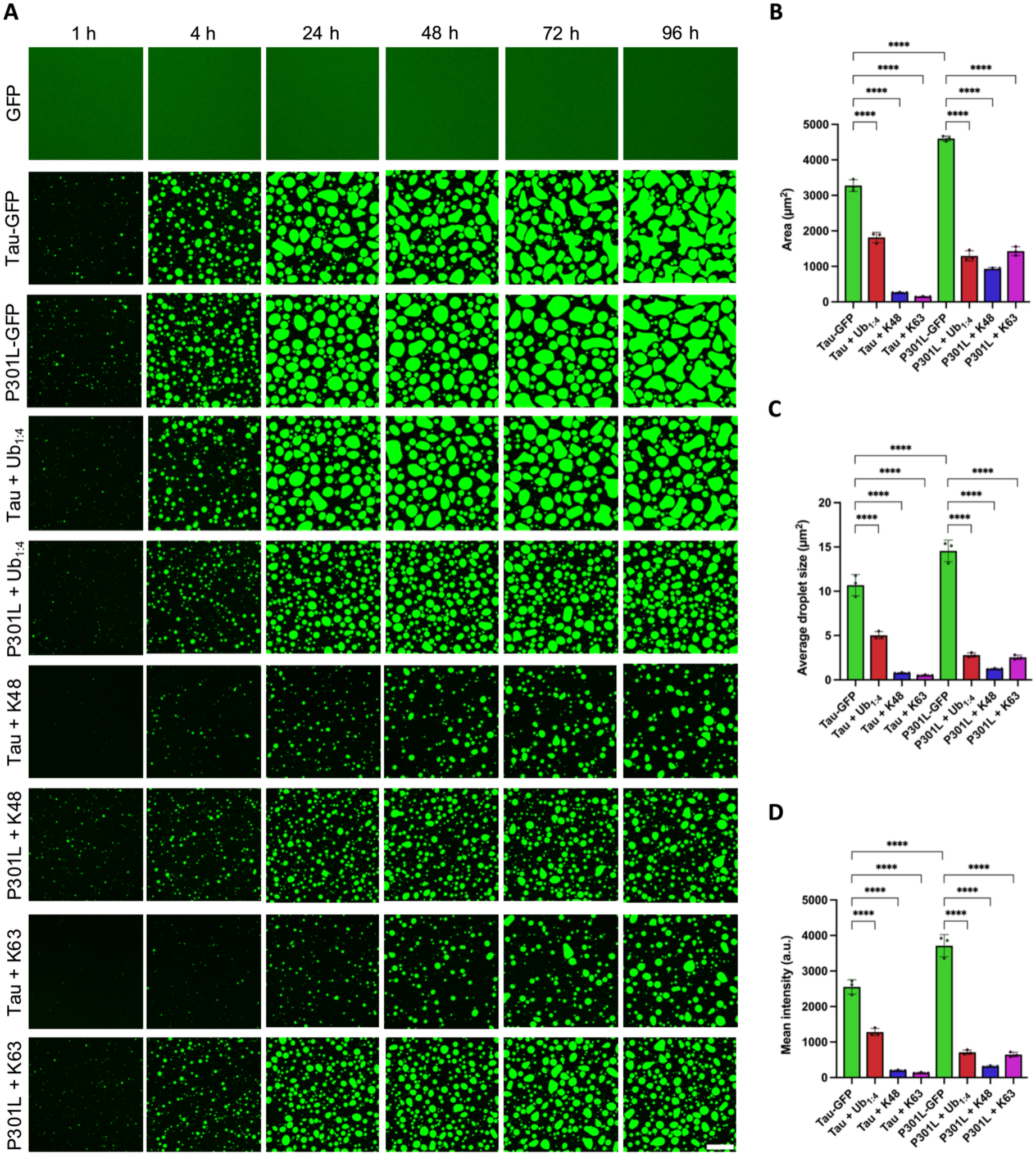
Ubiquitin differentially modulates Tau phase separation at physiological concentrations. **(A)** Tau-GFP and P301L-GFP readily undergo liquid-liquid phase separation at physiological concentrations (4 µM) under molecular crowding conditions (10% PEG), forming liquid-like condensates. Recombinant GFP did not phase separate under identical conditions, confirming that Tau mediates droplet formation. Representative droplets of Tau-GFP and P301L-GFP in the presence of four molar equivalents of mono-ubiquitin (Ub_1:4_) or two molar equivalents of di-ubiquitin chains (K48 or K63) are shown. All experiments were performed in LLPS buffer with 10% PEG. Scale bar, 25 µm. **(B–D)** Quantitative image analysis demonstrates that distinct ubiquitin species differentially regulate the phase separation behavior of both Tau-GFP and Tau-P301L-GFP. The extent of droplet formation **(B)**, average droplet size **(C),** and droplet fluorescence intensity **(D)** were determined after a 4-hour incubation with mono-ubiquitin at a 1:4 molar ratio (Ub_1:4_) or with K48- and K63-linked di-ubiquitin chains at a 1:2 molar ratio under crowding conditions. Graphs represent mean ± SD (one-way ANOVA with Holm–Sidak’s multiple comparisons test; n = 3 independent biological replicates; \*\*\*\**p* < 0.0001).

Consistent with previous reports (14), the P301L mutant exhibits accelerated phase separation, forming larger, more abundant condensates with higher fluorescence intensity than wild-type Tau **(Fig. 4B–D)**. In contrast, recombinant GFP alone fails to undergo LLPS under identical conditions, confirming that droplet formation is driven by Tau rather than the fluorescent tag **(Fig. 4A)**.

Adding mono-ubiquitin at a 1:4 Tau:ubiquitin molar ratio (Ub_1:4_) significantly reduced condensate abundance, average droplet size, and fluorescence intensity in both Tau variants **(Fig. 4B–D)**. The inhibitory effect is substantially greater for Tau-P301L, despite its enhanced propensity for phase separation. Fluorescently labeled ubiquitin partitions into Tau condensates, whereas ubiquitin alone does not undergo phase separation under identical conditions **(Supporting Fig. S7)**, indicating that ubiquitin is recruited through direct interaction with Tau. K48- and K63-linked di-ubiquitin at a 1:2 molar ratio produces a stronger suppression of LLPS than mono-ubiquitin **(Fig. 4B–D)**, consistent with the multivalent ubiquitin recognition mechanism identified by NMR and MD simulations.

K48- and K63-linked poly-ubiquitin chains differentially affected Tau phase separation. K63-linked chains inhibited wild-type Tau LLPS more effectively than K48-linked chains, whereas the opposite trend was observed for Tau-P301L **(Fig. 4B–D)**. Together with the NMR and MD analyses, these findings indicate that poly-ubiquitin-chain topology influences Tau recognition and that the P301L mutation alters the response of Tau condensates to poly-ubiquitin. These findings establish ubiquitin as a negative regulator of Tau phase separation and demonstrate that poly-ubiquitin exhibits enhanced inhibitory activity through multivalent engagement of the Tau repeat domain.

As an orthogonal quantitative assessment of microscopy observations, we also monitored Tau phase separation by microplate-based turbidity measurements at 300 nm and 600 nm (42). Mono-ubiquitin suppressed phase separation of both wild-type Tau and Tau-P301L in a concentration-dependent manner, with a more pronounced inhibitory effect observed for the P301L mutant **(Supporting Fig. S8A–H)**. These measurements independently confirmed the microscopy results, demonstrating suppression of Tau phase separation using both imaging- and turbidity-based approaches.

### Ubiquitin enhances the dynamics of Tau condensates

To determine whether ubiquitin influences the viscoelastic properties of Tau condensates, we monitored fluorescence recovery after photobleaching (FRAP) in Tau-GFP and Tau-P301L-GFP droplets. After 4 hours, condensates formed by both Tau variants exhibited highly dynamic, liquid-like behavior, with low immobile fractions and rapid fluorescence recovery **(Fig. 5A–C)**. Adding mono-ubiquitin (Ub_1:4_) produces only minor changes in immobile fractions but significantly accelerates fluorescence recovery in both Tau-GFP and Tau-P301L-GFP condensates, indicating enhanced molecular exchange between the droplets and the solvent (40).

**Figure 5.**
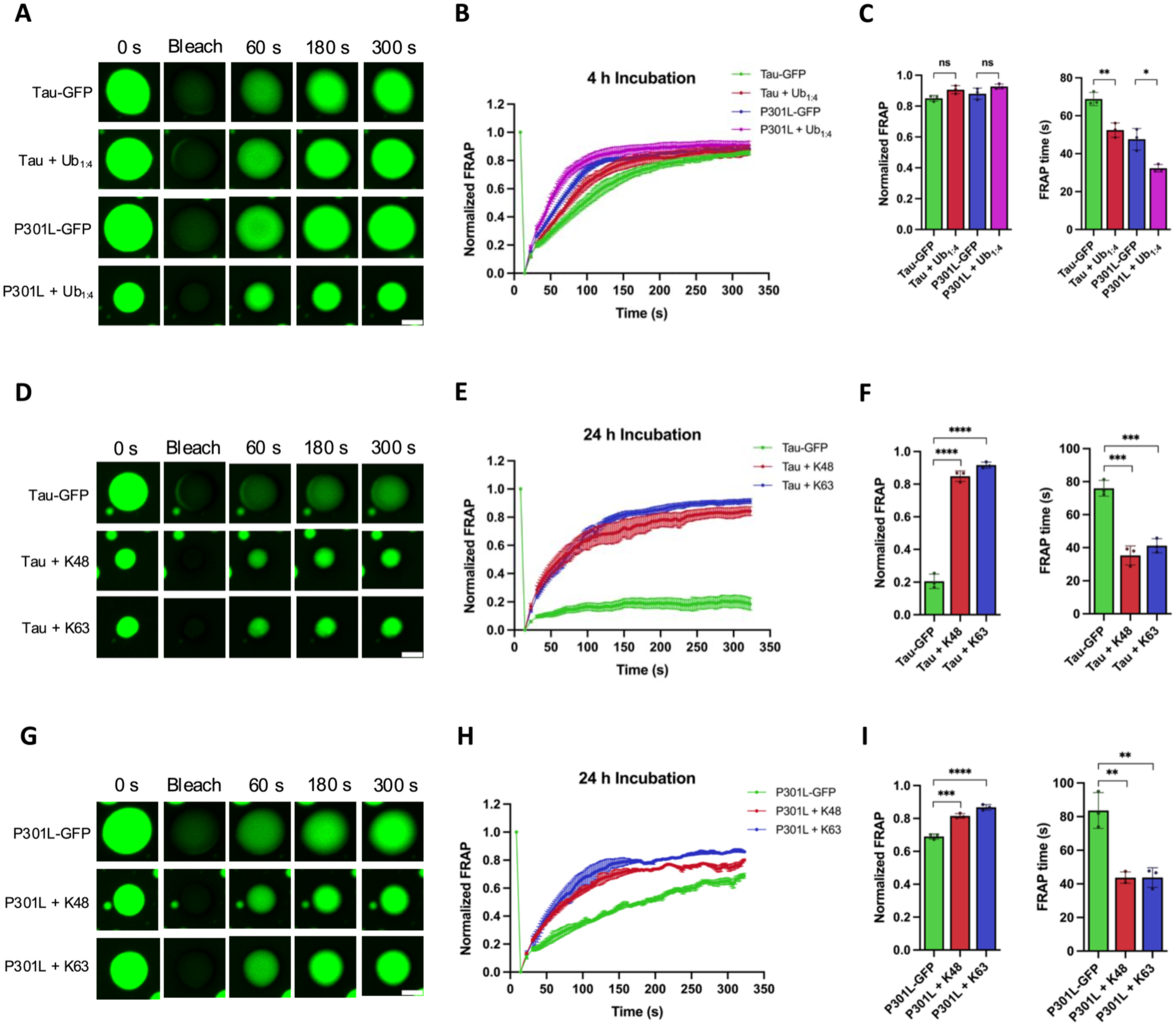
Ubiquitin and poly-ubiquitin chains enhance the dynamics of Tau liquid droplets. **(A–C)** Fluorescence recovery after photobleaching (FRAP) shows that mono-ubiquitin (mono-Ub) enhances the material dynamics of Tau condensates. Representative FRAP images of Tau-GFP and Tau-P301L-GFP condensates formed in the presence of four molar equivalents of mono-ubiquitin (Ub_1:4_) are shown **(A)**. Recovery curves demonstrate enhanced molecular exchange within both Tau-GFP and Tau-P301L-GFP condensates in the presence of mono-Ub **(B)**. Quantification of fluorescence recovery after 300 s and recovery half-times **(C)** confirms that mono-Ub significantly increases the dynamic properties of condensates formed by both Tau proteins. **(D–I)** K48- and K63-linked di-ubiquitin chains similarly enhance the material dynamics of both wild-type and P301L Tau condensates. Representative FRAP images of Tau-GFP **(D)** and Tau-P301L-GFP **(G)** condensates formed in the presence of two molar equivalents of K48- or K63-linked di-ubiquitin are shown. Corresponding FRAP curves **(E, H)** demonstrate increased molecular exchange within condensates relative to Tau alone, while quantification of fluorescence recovery after 300 s and recovery half-times **(F, I)** confirms that both ubiquitin chains significantly enhance the dynamic properties of condensates formed by wild-type and P301L Tau. FRAP experiments were performed after 4 h or 24 h incubation under crowding conditions (10% PEG) using 4 µM Tau-GFP or Tau-P301L-GFP in LLPS buffer. Graphs represent mean ± SD (One-way ANOVA with Holm–Sidak’s multiple comparisons test; n = 3 independent FRAP experiments; ns, not significant; \**p* < 0.05; \*\**p* < 0.01; \*\*\**p* < 0.001; \*\*\*\**p* < 0.0001).

After 24 hours of incubation, Tau condensates undergo substantial maturation and show markedly reduced molecular mobility **(Fig. 5D–I)**. Wild-type Tau droplets develop a large immobile fraction, whereas Tau-P301L droplets display irregular recovery profiles, consistent with structural heterogeneity and the emergence of aggregation-like states within the condensates. Adding either K48- or K63-linked poly-ubiquitin chains markedly restores condensate dynamics in both Tau variants, significantly increasing fluorescence recovery and shortening recovery half-times **(Fig. 5D–I)**. Although the irregular recovery pattern of Tau-P301L-GFP is not completely eliminated, both poly-ubiquitin chain types significantly improved its dynamic properties.

Together, these results demonstrate that ubiquitin influences not only Tau condensate formation but also condensate material properties following phase separation. Mono-ubiquitin enhances molecular exchange in newly formed condensates, whereas poly-ubiquitin promotes the dynamic behavior of more mature condensates. These functional effects are consistent with the multivalent Tau–ubiquitin interactions identified by NMR and MD simulations.

### Ubiquitin differentially modulates aggregation of wild-type and P301L Tau-MTBR

To determine whether the effects of ubiquitin extend beyond LLPS to later stages of Tau self-assembly, we examined heparin-induced aggregation of Tau-MTBR and the FTD-associated Tau-MTBR-P301L variant using Thioflavin T (ThT) fluorescence assays at 37 °C. Heparin was used as the standard *in vitro* polyanionic inducer, serving as a surrogate for heparan sulfate proteoglycans, which are the physiologically relevant polyanions in brain tissue (43).

Consistent with previous reports, Tau-MTBR-P301L aggregates more rapidly and reaches higher ThT fluorescence intensities than wild-type Tau-MTBR **(Fig. 6A)**, confirming the enhanced aggregation propensity of the disease-linked mutant. Adding mono-ubiquitin at a 1:4 molar ratio (Ub_1:4_) has little to no effect on the aggregation kinetics of wild-type Tau-MTBR **(Fig. 6B)**. In contrast, mono-ubiquitin (Ub_1:4_) increases ThT fluorescence during both the elongation and plateau phases of Tau-MTBR-P301L aggregation **(Fig. 6C)**. Direct comparison of both proteins in the presence of Ub_1:4_ shows a substantially greater effect on Tau-MTBR-P301L than on wild-type Tau **(Fig. 6D).**

**Figure 6.**
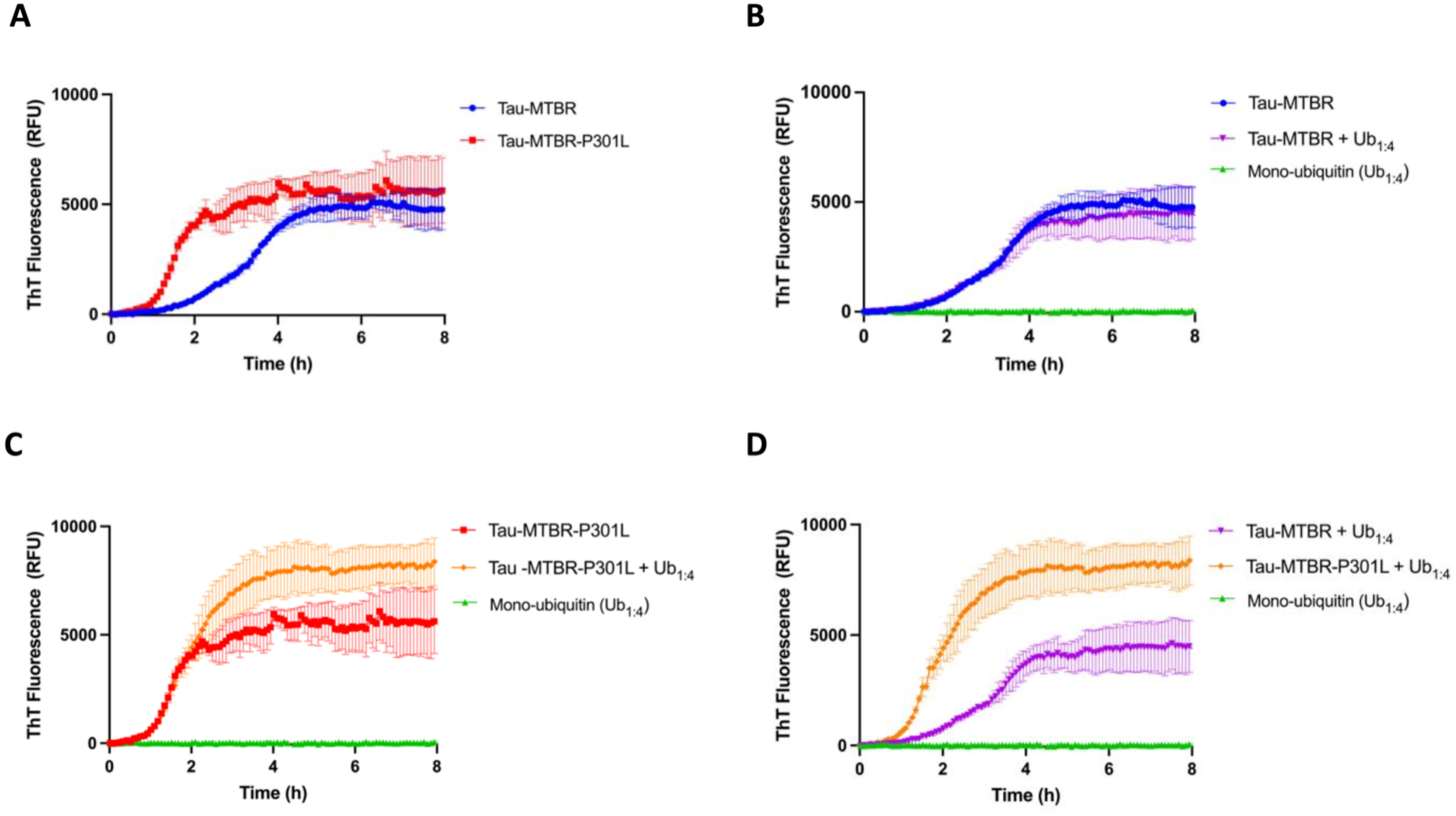
Mono-ubiquitin modulates Tau-MTBR aggregation independently of liquid-liquid phase separation. **(A–D)** Mono-ubiquitin directly modulates heparin-induced aggregation of Tau-MTBR independently of condensate formation. Aggregation of wild-type Tau-MTBR and the disease-associated variant Tau-MTBR-P301L (5 µM) was monitored by Thioflavin T (ThT) fluorescence in the presence of 3 kDa heparin (5 µM) at 37 °C **(A)**. The effect of four molar equivalents of mono-ubiquitin (Ub_1:4_) on aggregation kinetics was subsequently assessed for Tau-MTBR (**B**) and Tau-MTBR-P301L **(C)**. Direct comparison of both Tau variants in the presence of Ub_1:4_ **(D)** demonstrates that ubiquitin differentially modulates amyloid formation by the wild-type and mutant proteins. Graphs represent mean ± SD from three technical replicates.

To determine whether these aggregation effects are linked to condensate formation, we monitored turbidity (OD_600_) throughout the aggregation assays. Turbidity remains at baseline across all tested conditions **(Supporting Fig. S9)**, indicating no detectable LLPS during the aggregation reactions. Together, these findings demonstrate that mono-ubiquitin differentially modulates the aggregation of wild-type and P301L Tau-MTBR under conditions where detectable LLPS does not occur.

## Discussion

In this study, we identify Tau as a ubiquitin-binding protein and demonstrate that multivalent, non-covalent ubiquitin recognition regulates multiple stages of Tau self-assembly. Using biochemical, biophysical, structural, and computational approaches, we show that Tau engages ubiquitin through four distinct binding elements distributed across its microtubule-binding repeat region (MTBR). These interactions modulate condensate formation and dynamics and differentially influence the amyloid aggregation of wild-type and disease-associated Tau variants.

Strikingly, three of these ubiquitin-binding elements within Tau’s MTBR are positioned adjacent to established amyloidogenic motifs. The R2 element is located adjacent to the PHF6* motif, whereas the R3 element is located adjacent to the major Tau aggregation motif PHF6 (44, 45), separated by only two residues. Similarly, the R4 element is located adjacent to the recently described PAM4 motif (39). By contrast, the R1 element represents a previously unrecognized ubiquitin-binding site within a region implicated in other functionally relevant Tau interactions, including microtubule binding and Tau self-association (46). Thus, ubiquitin recognition spans the MTBR, with three binding elements positioned near amyloid-promoting regions of Tau. This spatial distribution suggests a potential mechanistic link between non-covalent ubiquitin recognition and the intermolecular interactions governing Tau self-assembly, whereby ubiquitin binding could compete with or remodel contacts that promote condensate formation and amyloid assembly.

Our structural and biophysical analyses support a model in which these sites cooperate to mediate multivalent ubiquitin recognition. NMR spectroscopy, MD simulations, co-evolution analysis, and AlphaFold-Multimer predictions converged on a common interaction mechanism in which multiple weak Tau–ubiquitin contacts cooperate to generate higher-avidity recognition (47, 48). Because multivalency is a fundamental determinant of both condensate formation and maturation (10), engagement of multiple ubiquitin-binding sites along the MTBR provides a plausible mechanism by which ubiquitin could remodel the interaction network underlying Tau self-assembly. This model also provides a structural explanation for Tau’s enhanced interaction with mono-ubiquitin and K63-linked poly-ubiquitin species relative to K48-linked chains. K63-linked poly-ubiquitin displays extended chain architectures that enable simultaneous engagement of multiple ubiquitin units, whereas the more compact conformational ensembles of K48-linked chains may restrict access to their hydrophobic interaction surfaces (40, 49). Altogether, these observations suggest that ubiquitin-chain topology may provide an additional layer of regulation of Tau assembly.

The functional consequences of Tau–ubiquitin interactions are most apparent in Tau phase behavior. Both mono- and poly-ubiquitin inhibited condensate formation while enhancing the liquid-like properties of the remaining condensates. These observations suggest that ubiquitin regulates Tau condensates at two related but distinct levels: limiting condensate assembly and opposing condensate maturation toward less dynamic, gel-like or solid-like states. A plausible mechanism is that engagement of multiple MTBR sites reduces the effective interaction valency available for Tau–Tau association, thereby disrupting the intermolecular contacts that drive condensate formation and maturation. A conceptually similar mechanism was demonstrated for UBQLN2, in which ubiquitin binding disrupts the multivalent interactions required for phase separation (50).

More broadly, our findings support emerging models in which PTMs and regulatory binding partners control biomolecular condensates by tuning interaction valency and network connectivity (17, 18). The multivalent ubiquitin-recognition mechanism identified in this study therefore provides a direct link between the structural and functional effects of ubiquitin on Tau self-assembly.

The effects of ubiquitin on Tau condensates may have important implications for tauopathy, as the material state of these condensates can influence their propensity to undergo pathological maturation. Previous studies showed that prolonged incubation of Tau condensates promotes the transition of initially liquid-like condensates toward less dynamic states associated with amyloid aggregation (8, 9). By suppressing condensate formation and enhancing the liquid-like behavior of both wild-type and mutant Tau condensates that do form, non-covalent ubiquitin binding may oppose early steps of pathological condensate maturation. This interpretation expands the conventional view of ubiquitin enrichment within pathological protein aggregates during neurodegeneration (51). Rather than acting solely as a marker of impaired proteostasis and protein degradation, ubiquitin may directly regulate the interaction networks that precede aggregate formation.

Importantly, the effects of ubiquitin were not uniformly protective across the Tau self-assembly pathway. Under heparin-induced aggregation conditions, in which no detectable LLPS occurred, mono-ubiquitin had little effect on wild-type Tau-MTBR, but consistently enhanced ThT fluorescence during the elongation and plateau phases for the disease-associated Tau-MTBR-P301L variant. These findings indicate that the consequences of non-covalent ubiquitin recognition depend on Tau’s molecular context and cannot be inferred solely from its effects on phase separation. Because the P301L mutation lies within the MTBR and alters both the phase separation and aggregation propensity of Tau, ubiquitin engagement may yield distinct outcomes depending on the assembly state of the protein. We note that ubiquitin affinity was determined only for wild-type Tau, and future studies should address whether the P301L variant displays altered binding affinity to mono- and poly-ubiquitin chains.

The apparent affinity we measure (*K*_D_ ≈ 220–260 µM) is low relative to the bulk cellular free ubiquitin pool, which has been quantified at roughly 20 µM in HEK293 cells, although the free, unconjugated form constitutes the majority of total ubiquitin in brain tissue (≈60% in mouse brain and ≈82% in human frontal cortex) (52). Non-covalent Tau–ubiquitin engagement is therefore unlikely to arise from saturation of individual binding sites in bulk cytoplasm and is instead expected to depend on avidity from multivalent engagement of poly-ubiquitin chains or ubiquitinated substrates, and on local enrichment of ubiquitin in the compartments where Tau assembles. Consistent with this, mCherry-tagged ubiquitin partitions into Tau condensates in our assays **(Supporting Fig. S7)**, and ubiquitin is enriched within pathological inclusions in neurodegenerative disease (51), environments in which the local ubiquitin concentration may considerably exceed the bulk cytoplasmic value. Notably, the effects reported here occur at ubiquitin concentrations of 8–20 µM, within the range of the cellular free pool and well below the apparent *K*_D_, which is consistent with avidity-driven, sub-saturating engagement, although a contribution from the crowded assay conditions cannot be excluded.

Altogether, whether ubiquitin binding is ultimately protective or pathogenicity-enhancing will likely depend on multiple factors, including ubiquitin and protein concentrations, ubiquitin chain architecture, other interaction proteinogenic and non-proteinogenic partners (like heparin), and Tau modification state. Whether this principle extends beyond Tau to other aggregation-prone ubiquitin-binding proteins associated with neurodegenerative disease will be an important question for future studies.

Although this work primarily examined interactions between Tau and free ubiquitin species *in vitro*, several key questions remain. Disease-relevant phosphorylated Tau can undergo phase separation without molecular crowders (53), raising the question of whether phosphorylation alters non-covalent Tau–ubiquitin recognition or condensate formation under more physiologically relevant conditions. The overlap between ubiquitin-binding elements and aggregation-prone regions of the MTBR further suggests a more complex interplay between ubiquitin recognition and disease-linked Tau modifications. Importantly, whether Tau–ubiquitin interactions also serve physiological functions remains unknown. Histone deacetylase 6 (HDAC6) provides an interesting precedent, interacting with Tau through the MTBR while independently recognizing ubiquitin during cellular protein quality control (54, 55).

Because the MTBR also mediates both Tau self-association and microtubule binding, ubiquitin binding could influence Tau–microtubule association, interactions with other binding partners, or Tau conformational dynamics. Future studies should therefore address these possibilities alongside the effects of Tau modifications, ubiquitin-chain topology, covalent ubiquitination, and cellular protein quality control pathways. Extending these observations to cellular and disease-relevant models will be essential for defining the roles of non-covalent ubiquitin recognition in physiological Tau regulation and pathological assembly.

Collectively, our findings identify Tau as a ubiquitin-binding protein that engages ubiquitin through four binding elements distributed along the MTBR. These interactions enable multivalent, non-covalent ubiquitin recognition that modulates Tau condensate formation and material properties while differentially regulating amyloid assembly depending on the molecular context of Tau. Together, our results establish non-covalent ubiquitin recognition as a previously unrecognized mechanism that regulates multiple stages of Tau self-assembly. More broadly, these findings expand the role of ubiquitin beyond protein quality control and suggest that ubiquitin can directly influence the transition of Tau from a soluble protein to pathological assemblies. Beyond ubiquitin-mediated degradation, non-covalent ubiquitin recognition may provide an additional proteostatic mechanism by limiting condensate formation and maintaining the dynamic properties of Tau assemblies. However, its consequences for subsequent amyloid formation depend on the molecular context of Tau. This framework places ubiquitin at the interface between proteostasis and protein self-assembly and identifies multivalent ubiquitin recognition as a potential mechanistic link between these processes in tauopathy.

## Materials and Methods

### ALS inclusion body protein network analysis

The ALS inclusion body proteome (52 proteins) was derived from published proteomic datasets (33). Protein–protein interaction networks were constructed in Cytoscape v3.8 (56) using interactions retrieved via the PSICQUIC interface (57). Network significance was assessed by comparison with 10,000 random protein networks of the same size, sampled from the human BioGRID interactome using a custom Python workflow.

### Proteome overlap analysis of ALS inclusion body proteins

The overlap among the ALS inclusion body proteome (33), the stress granule proteome (34), and the human ubiquitin interactome retrieved from BioGRID (35) was assessed using Venn diagram analysis (Bioinformatics & Evolutionary Genomics, Ghent University).

### Structural modeling of Tau–ubiquitin complexes

Ubiquitin structural models were based on the crystal structure of human ubiquitin (PDB: 1D3Z) (58). A Tau-MTBR peptide (residues 308–319; PDB: 5N5B) was docked onto ubiquitin using ClusPro2 (59). Evolutionary coupling analysis was performed using the ComplexContact web server (60) via RaptorX to assess support for the predicted interaction interface. Full-length Tau (UniProt: P10636-8) and ubiquitin (UniProt: P0CG47) sequences were submitted with default parameters. Homologous sequences were identified using HHblits, and a paired multiple sequence alignment was constructed for deep learning-based residue coevolution analysis. The resulting residue contact probability maps were compared with the predicted docking interface, with higher covariation scores shown in darker shades. Additional Tau–ubiquitin complexes were generated using AlphaFold2-Multimer v3 (61) implemented in ColabFold (62).

### Radiolabeled ubiquitin pull-down assays

Candidate ubiquitin-binding proteins were cloned into the mammalian expression vector pcDNA3.1(+) and synthesized by *in vitro* transcription/translation using the TNT T7 Coupled Reticulocyte Lysate System (Promega) according to the manufacturer’s instructions. Radiolabeled proteins were synthesized using the TNT T7 (Promega; L4610) according to the manufacturer’s instructions. Open reading frames encoding the 11 candidate proteins were cloned into a pcDNA3.1(+) mammalian expression vector containing a T7 promoter for cell-free transcription. *In vitro* transcription/translation reactions were supplemented with T7 RNA polymerase (ThermoFisher Scientific), RiboLock RNase inhibitor (Thermo Scientific), an amino acid mixture lacking methionine, 1 µg plasmid DNA, and EasyTag^TM^ L-[^35^S]-methionine (PerkinElmer), and incubated for 90 min at 30 °C. Lysates containing specific radiolabeled proteins were incubated with His-tagged mono-ubiquitin (Mono-Ub), K48-linked di-ubiquitin (polyUb-K48), or K63-linked di-ubiquitin (polyUb-K63) (UBPBio) in binding buffer (50 mM Tris-HCl, 150 mM NaCl, 1% Nonidet P-40, 5% glycerol, cOmplete™ protease inhibitor cocktail, pH 7.4) for 3 hours at 4 °C with gentle mixing. Complexes were captured using nickel affinity beads equilibrated with binding buffer, then centrifuged, washed extensively with binding buffer and cold PBS, and finally analyzed by SDS-PAGE. Ubiquitin probes were visualized by Coomassie staining, whereas radiolabeled proteins were detected by phosphorimaging. All experiments were performed at least in three independent biological replicates.

### Protein expression and purification

C-terminally Avi-tagged Tau441 (Tau-hT40; UniProt P10636-8) was cloned into the tandem-tag pSUMO expression vector described in (63). The construct was expressed in *Escherichia coli* AVB101 cells grown in TYH medium (64) supplemented with 10 µM biotin, 10 µg/mL chloramphenicol, and 50 µg/mL kanamycin, then induced with 1.5 mM IPTG for 3 h at 37 °C. Biotinylated Tau-Avi was purified using a previously described tandem-tag strategy involving nickel-affinity and heparin chromatography, followed by TEV cleavage and removal of the affinity tags (63).

Full-length Tau441 (Tau-hT40), Tau-P301L, Tau-MTBR, and Tau-MTBR-P301L were cloned into the pT7NT* (MaSp) expression vector, which contains an N-terminal His_6_-tagged NT* solubility tag (65) and a TEV cleavage site, as described previously (66). Recombinant proteins were expressed in *E. coli* BL21 STAR™(DE3) cells and purified using the NT*-based strategy (66).

Tau-GFP, Tau-P301L-GFP, and GFP expression constructs were kindly provided by Prof. Nicholas M. Kanaan (Michigan State University). Proteins were expressed in *E. coli* T7 Express cells and purified by nickel-affinity, size-exclusion, and anion-exchange chromatography, as described previously (67).

Human mono-ubiquitin and mCherry-tagged ubiquitin were custom synthesized by GenScript in pET22b(+) vectors. Proteins were expressed in *E. coli* BL21 STAR™(DE3) cells grown in LB and TB media, respectively, after induction with 1 mM IPTG for 4 hours at 37 °C. Proteins were purified by nickel-affinity followed by size-exclusion chromatography. The column was equilibrated in 20 mM Tris-HCl, 150 mM NaCl, 5% glycerol, and 1 mM TCEP (pH 7.4).

### Biolayer interferometry

Biolayer interferometry (BLI) measurements were performed on an OCTET RED96 instrument (ForteBio). 5 µg/mL biotinylated Tau-Avi was immobilized on Super-Streptavidin biosensors (ForteBio), and mono-ubiquitin was titrated from 4 mM in BLI Kinetics buffer (50 mM Tris, 150 mM NaCl, 0.5 mM TCEP, 0.05% Tween-20, pH 7.5). Measurements were performed at 25 °C with orbital shaking at 1000 rpm. Association and dissociation curves were analyzed using Octet Data Analysis software v9.0 to determine equilibrium dissociation constants (K_D_). All experiments were performed in triplicate.

### Microscale thermophoresis

Tau proteins were labeled using the MO-L011 RED-NHS 2^nd^ Generation protein labeling kit (NanoTemper). Ubiquitin binding was assessed on a Monolith™ NT.115 instrument (NanoTemper). Mono-ubiquitin was titrated against 200 mM labeled Tau in MST buffer (50 mM Tris-HCl, 150 mM NaCl, 10 mM MgCl_2_, and 0.05% Tween-20, pH 7.4). Measurements were acquired at 80% MST laser power (5 s off, 20 s on, 5 s off). Binding curves were fitted using MO Affinity Analysis software to determine K_D_ values. All experiments were performed in triplicate.

### Nuclear magnetic resonance spectroscopy

Uniformly ^15^N-labeled Mono-Ub and uniformly ^13^C, ^15^N-labeled Tau-MTBR were produced by expression in M9 minimal medium using standard isotopic labeling protocols (68). The NMR samples contained 0.2 mM of U-[^13^C,^15^N] Tau-MTBR or U-[^15^N] Mono-Ub in 20 mM Tris-HCl pH 6.5, 100 mM NaCl, and 6% D_2_O for the lock. NMR spectra were acquired at 298 K on a Bruker Avance III HD 800 MHz spectrometer equipped with a cryogenic probe. Data were processed in TopSpin 3.7 (Bruker) or NMRPipe (69) and analyzed by CCPNMR (70). Backbone resonance assignments of Tau-MTBR were taken from our previous work (66), whereas mono-ubiquitin assignments obtained from (71) were verified using three-dimensional ^15^N-edited NOESY-HSQC and TOCSY-HSQC experiments. Protein-protein interactions were analyzed by monitoring chemical shift perturbations in two-dimensional [^1^H,^15^N] HSQC spectra during titration of unlabeled protein into its corresponding labeled binding partner.

Average backbone amide chemical shift perturbations (Δδ_avg_) were calculated as Δδ_avg_ = (Δδ_N_^2^/50 + Δδ_H_^2^/2)^0.5^, where Δδ_N_ and Δδ_H_ are the chemical shift changes of the backbone amide nitrogens and protons, respectively. Titration curves obtained for U-[^15^N] Mono-Ub with unlabeled Tau-MTBR were analyzed using a two-parameter nonlinear least-squares fit with a one-site binding model corrected for dilution effects, as described previously (72), Equation 1:

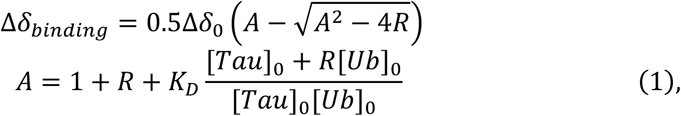

where Δδ_binding_ is the chemical shift perturbation at a given protein/protein ratio; Δδ_0_ is the chemical shift perturbation at 100% Mono-Ub bound; R is the [Tau-MTBR]/[Mono-Ub] ratio at a given point; [Ub]_0_ and [Tau]_0_ are the concentrations of the starting U-[^15^N] Mono-Ub sample and the Tau-MTBR titrant stock solution, respectively; and *K*_D_ is the dissociation constant.

### Molecular dynamics simulations

Starting structures were generated from PDB 6UD0 for head-to-tail di-ubiquitin and PDB 1A5R for SUMO1. Tau-F (UniProt P10636-8; residues K225–K375, encompassing R1–R4 and flanking regions) was predicted using AlphaFold2 (61), whereas the α-synuclein structure was obtained from the Protein Ensemble Database (PED00024e001) (73). The Tau–di-ubiquitin complex was built by directed docking using the HADDOCK 2.4 web server (74), with Tau residues ^281^KLDLSN^286^ (Patch 1) and ^314^DLSKVT^319^ (Patch 2), and ubiquitin interface residues K6, ^43^LIFAG^47^, Q49, and ^67^HLVLR^71^. Corresponding residues in the second ubiquitin moiety (K82, ^119^LIFAG^123^, Q125, and ^143^HLVLR^147^) were extrapolated from mono-ubiquitin NMR titrations. Thus, although the canonical hydrophobic patch is conserved across the ubiquitin fold, simultaneous bivalent engagement was not directly determined by NMR and should be regarded as a working model. The α-synuclein–SUMO1 complex was similarly built by directed docking using the active residues reported by Panigrahi et al. (41): α-synuclein residues G37–V53 and SUMO1 residues H22–V27, K33–V50, and E54.

Simulations were performed on the Tau-MTBR R1–R4 (residues K225–K375, 151 aa) in complex with di-ubiquitin and for the α-synuclein–SUMO1 complex. The highest-ranked docking models were subjected to all-atom molecular dynamics (MD) simulations using GROMACS v. 2021.5 (75), the CHARMM36m force field (76), TIP3P water model, and 0.15 M KCl.

Simulations used a 2-fs time step with the Verlet leapfrog algorithm and a 1.2-nm cutoff for short-range electrostatic and Lennard-Jones interactions. Long-range electrostatic interactions were treated with the Particle Mesh Ewald method (77). Solute and solvent were separately coupled to 310 K using a modified Berendsen thermostat (V-rescale; τ = 0.1 ps) (78), and pressure coupling was fixed at 1 bar using a C-rescale barostat (79) with an isothermal compressibility of 4.5 × 10^−5^ bar^−1^. Periodic boundary conditions were applied in all three dimensions. After energy minimization, systems underwent 1 ns of NVT and 1 ns of NPT equilibration with position restraints on protein heavy atoms, followed by unrestrained NPT production simulations at 310 K and 1 bar.

Two independent simulations were performed for each system. For the Tau R1–R4/di-ubiquitin, replicas were set to 540 ns; replica 1 was terminated at 440 ns after complete dissociation at approximately 270 ns, whereas replica 2 remained associated throughout the full 540 ns. For α-synuclein–SUMO1, replicas were set to 400 ns, extending the 300-ns simulations reported by Panigrahi et al. (41); replica 1 remained associated throughout 400 ns, whereas replica 2 was terminated at approximately 90 ns following complete dissociation.

Residue-residue contacts were analyzed throughout the trajectories using bound complexes: replica 2 for Tau R1–R4/di-ubiquitin and replica 1 for α-synuclein–SUMO1. Residues were considered in contact when their minimum inter-residue distance was <7.4 Å, encompassing the largest cutoff used in the RING relaxed mode (80) and capturing hydrogen bonds, salt bridges, π–cation, π–π, and van der Waals interactions. Contact occupancy was calculated as the percentage of simulation frames containing each residue pair and visualized using a square-root scale (power-law, γ = 0.5) to retain low-occupancy contacts. Color-bar labels indicate the corresponding occupancy values. Trajectories were analyzed and visualized using Visual Molecular Dynamics (81) and ChimeraX (82).

### Induction of liquid-liquid phase separation

GFP-tagged Tau proteins were diluted in LLPS buffer (10 mM HEPES, 150 mM NaCl, 0.1 mM EDTA, 2 mM DTT, pH 7.4) to a final protein concentration of 4 µM unless otherwise indicated. Mono-ubiquitin and di-ubiquitin (K48 and K63) were added to final concentrations of 16 µM and 8 µM, respectively. LLPS was induced at room temperature by adding 10% PEG 3000 before imaging or turbidity measurements. Recombinant GFP served as a negative control.

### Confocal microscopy and fluorescence recovery after photobleaching

Confocal microscopy was performed on a Nikon A1+ laser-scanning confocal microscope equipped with a 60× oil-immersion objective (NA 1.4). The system was controlled using Nikon Elements Imaging Software (v5). All samples were imaged under identical settings. Droplet area, average droplet size, and fluorescence intensity were quantified in Fiji (ImageJ), using a fixed intensity threshold (360–4095 gray values) to remove background fluorescence.

Fluorescence recovery after photobleaching (FRAP) was performed after 4 h and 24 h of incubation. Tau condensates were photobleached at 50% laser power. Images were acquired before, during, and after bleaching over a 5 min recovery period at 2 s intervals. Fluorescence recovery curves were normalized to the reference and background signals, and half-times of recovery and immobile fractions were calculated using Nikon NIS-Elements Software (v5). Each experiment was performed in three independent replicates.

### Tau colocalization assays with mCherry-Ub

Tau colocalization assays were performed on a Leica DMi8 inverted fluorescence microscope (Leica Microsystems) equipped with GFP and mCherry filters, a DFC7000 GT camera, and an HC PL FLUOTAR 100x/1.32 Oil PH3 objective. Samples were prepared as described above, and mCherry-tagged ubiquitin (mCherry-Ub) was added before PEG-induced phase separation. Images were acquired sequentially using identical microscope settings.

### Turbidity Measurements

Tau proteins were diluted to 10 µM in LLPS buffer supplemented with 40 µM ZnCl_2_. To assess the concentration-dependent effects of mono-ubiquitin, Tau was pre-incubated with increasing concentrations of mono-ubiquitin, and LLPS was induced by adding 10% PEG 3000 (resulting in Tau concentration of 5 µM). Turbidity was monitored by measuring optical density at 300 nm and 600 nm every 2 minutes using a Synergy HTZ Multi-Mode Microplate Reader (BioTek) at 25 °C with slow orbital shaking. Measurements were performed in four technical replicates, and buffer controls were subtracted for baseline correction.

### Thioflavin T Fluorescence Assays

Aggregation of wild-type and P301L Tau-MTBR was monitored by Thioflavin T (ThT) fluorescence. Proteins were diluted to 5 µM in aggregation buffer (20 mM Tris-HCl, 50 mM NaCl, 1 mM TCEP, 1 mM EDTA, pH 7.4) containing 50 µM ThT and, where indicated, 20 µM mono-ubiquitin. Aggregation was initiated by adding 5 µM unfractionated heparin and monitored at 37°C on the plate reader. ThT fluorescence was recorded every 5 min using excitation and emission wavelengths of 450 and 490 nm, respectively. Measurements were performed in triplicate, and buffer-subtracted fluorescence traces were used for subsequent analyses.

### Statistical analysis

Statistical analyses were performed in GraphPad Prism unless otherwise indicated. Differences among experimental groups were assessed using one-way ANOVA with the Holm–Sidak or Sidak multiple-comparison tests. BLI, MST, and NMR binding data were analyzed by nonlinear regression using the corresponding binding models. NMR spectra were processed and analyzed using CCPNMR (64). Statistical significance was defined as *P* ≤0.05. No data points or experimental replicates were excluded from the analyses.

## Acknowledgments

We thank members of the Kanaan laboratory for producing and purifying the recombinant GFP-tagged Tau proteins used in this study. We also thank Tessa Grabinski for preparing the plasmids encoding the Tau domain constructs used in this study.

## Funding

K.M (11D0122N) and L.F.D.A (1163625N) were funded by PhD fellowships for Fundamental Research from the Research Foundation - Flanders (Fonds Wetenschappelijk Onderzoek; FWO). T.L. was funded by a Postdoctoral Innovation Mandate (HBC.2022.0194) awarded by the Flanders Innovation & Entrepreneurship Agency (Agentschap Innoveren & Ondernemen; VLAIO).

G.J.G. was funded by the VUB Interdisciplinary Research Program for Excellence on Cancer Research and the FWO G0C7514N grant. P.T. was funded by grants K124670 and K131702 from the National Research, Development and Innovation Office (NKFIH, Hungary). This work used computational resources provided by UNC Supercómputo (CCAD), Universidad Nacional de Córdoba, Argentina.

## Author Contributions

**Kevin Muwonge:** Writing – review & editing, Writing – original draft, Visualization, Methodology, Investigation, Formal analysis, Funding acquisition. **Luis Fernando Durán-Armenta:** Writing – review & editing, Writing – original draft, Visualization, Methodology, Investigation, Formal analysis, Funding acquisition. **Tamas Lazar:** Investigation, Methodology, Formal analysis, Validation. **Attila Mészáros:** Methodology, Investigation, Validation. **Alexander Volkov:** Investigation, Formal analysis, Validation. **Alvaro Navarro:** Investigation, Visualization. **Maria-Laura Fernández:** Investigation, Visualization. **Cristina Marino-Buslje:** Validation, Supervision. **Gustavo J. Gutierrez:** Methodology, Investigation, Validation, Supervision. **Nicholas M. Kanaan:** Methodology, Validation, Supervision. **Remy Loris:** Writing – review & editing, Supervision, Conceptualization, Funding acquisition. **Peter Tompa:** Writing – review & editing, Supervision, Conceptualization, Funding acquisition, Project administration.

## Declaration of interests

The authors declare they have no known competing financial interests or personal relationships that could have appeared to influence the work reported in this paper.

## Data Availability

Data will be available upon reasonable request to the corresponding authors.

## Notes

### Competing Interest Statement

The authors have declared no competing interest.

### Summary of Updates

This version of the manuscript has been revised to improve clarity in the results section. Specifically, Ub4 has been updated to Ub1:4 to indicate the molar ratio more clearly. Typos have been corrected.

