## Supporting Information for "Multivalent ubiquitin recognition regulates multiple stages of Tau self-assembly"

### Supporting Information for
### Multivalent ubiquitin recognition regulates multiple stages of Tau self-assembly

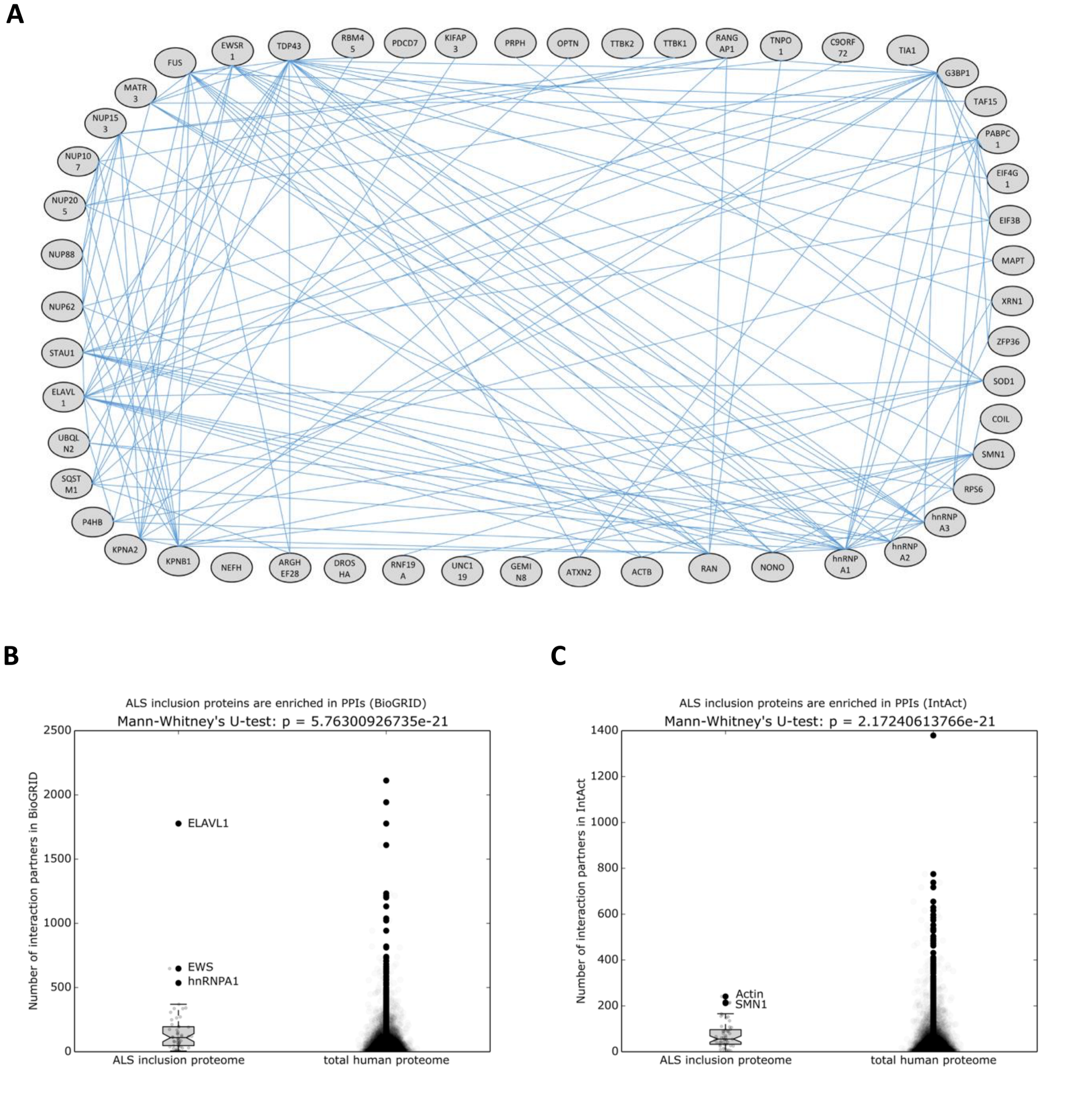


**Figure S1. ALS inclusion body proteins form a highly interconnected protein–protein interaction network.**

**(A–C)** Network analysis shows that the ALS inclusion body proteome forms a dense, interconnected protein–protein interaction network relative to the human interactome. The protein–protein interaction network of the 52-protein ALS inclusion body proteome is shown **(A)**. Comparisons with randomly sampled human protein networks of identical size, generated from the BioGRID **(B)** and IntAct **(C)** interactomes, show that ALS inclusion body proteins exhibit significantly greater network connectivity than expected by chance, supporting a densely interconnected disease-associated protein interaction network.

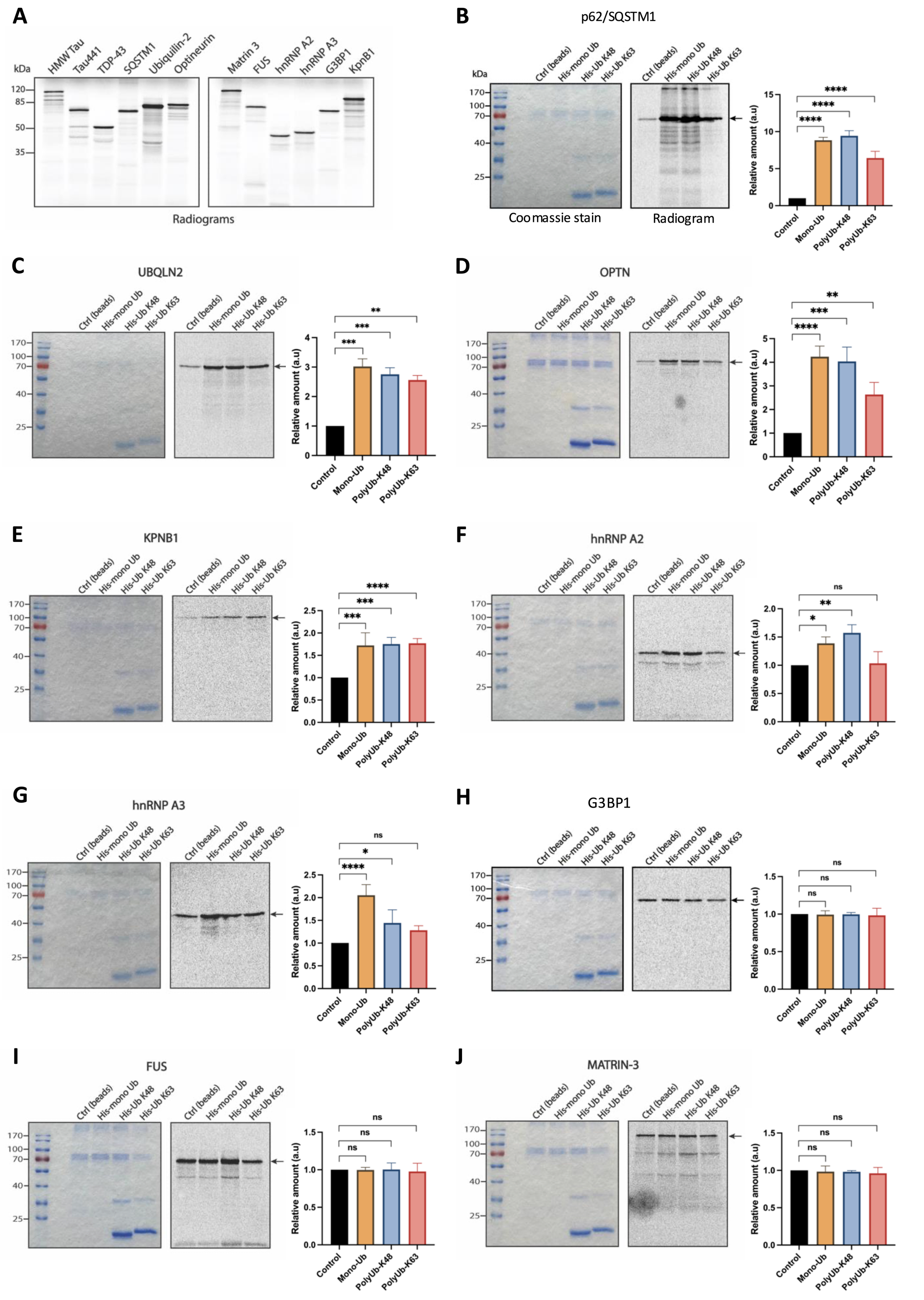


**Figure S2. Experimental validation of predicted ubiquitin-binding proteins by pull-down assays.** **(A–J)** Candidate ubiquitin-binding proteins (UBPs) identified by the integrative network analysis shown in Figure 1 were experimentally evaluated for ubiquitin binding. Radiolabeled candidate UBPs were synthesized using an *in*

*vitro* cell-free expression system and verified by SDS–PAGE followed by autoradiography **(A)**. **(B–J)** Ubiquitin pull-down assays classified the predicted UBP candidates as strong, weak, or non-ubiquitin binders, thereby benchmarking the screening strategy against established ubiquitin-binding proteins. Previously characterized ubiquitin receptors, including p62, Ubiquitin-2, and Optineurin, exhibited the strongest ubiquitin binding, validating the assay **(B–D)**. Tau displayed similarly robust ubiquitin-binding activity despite not being previously recognized as a ubiquitin-binding protein, whereas other candidates exhibited either weak or no detectable ubiquitin binding **(E–J)**. Graphs represent mean ± SD (One-way ANOVA with Sidak's multiple comparison test; ns, not significant; *$p < 0.05$; **$p < 0.01$; ***$p < 0.001$; ****$p < 0.0001$; n = 3 independent biological replicates).

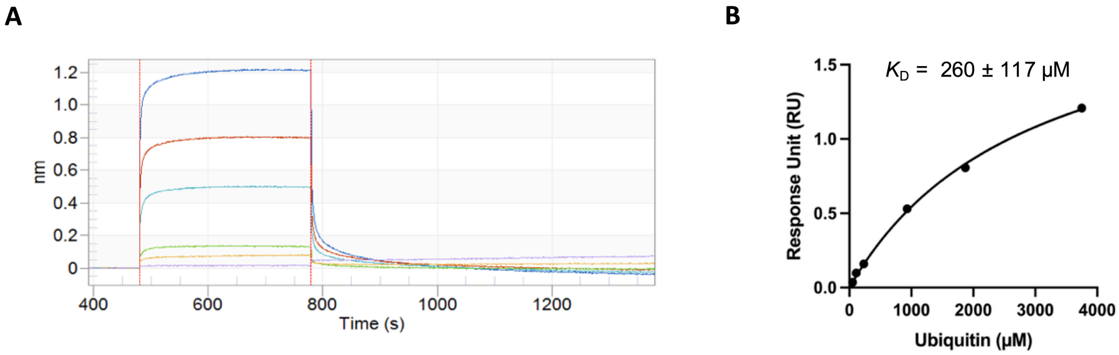


**Figure S3. Biolayer interferometry analysis of Tau–ubiquitin interaction**

**(A)** Representative biolayer interferometry (BLI) sensorgrams showing concentration-dependent binding of mono-ubiquitin to biotinylated Avi-tagged Tau441 immobilized on Super-streptavidin (SSA) biosensors. **(B)** Equilibrium binding responses plotted as a function of ubiquitin concentration and fitted to a 1:1 binding model, yielding a dissociation constant ($K_D$) of 260 ± 117 µM.

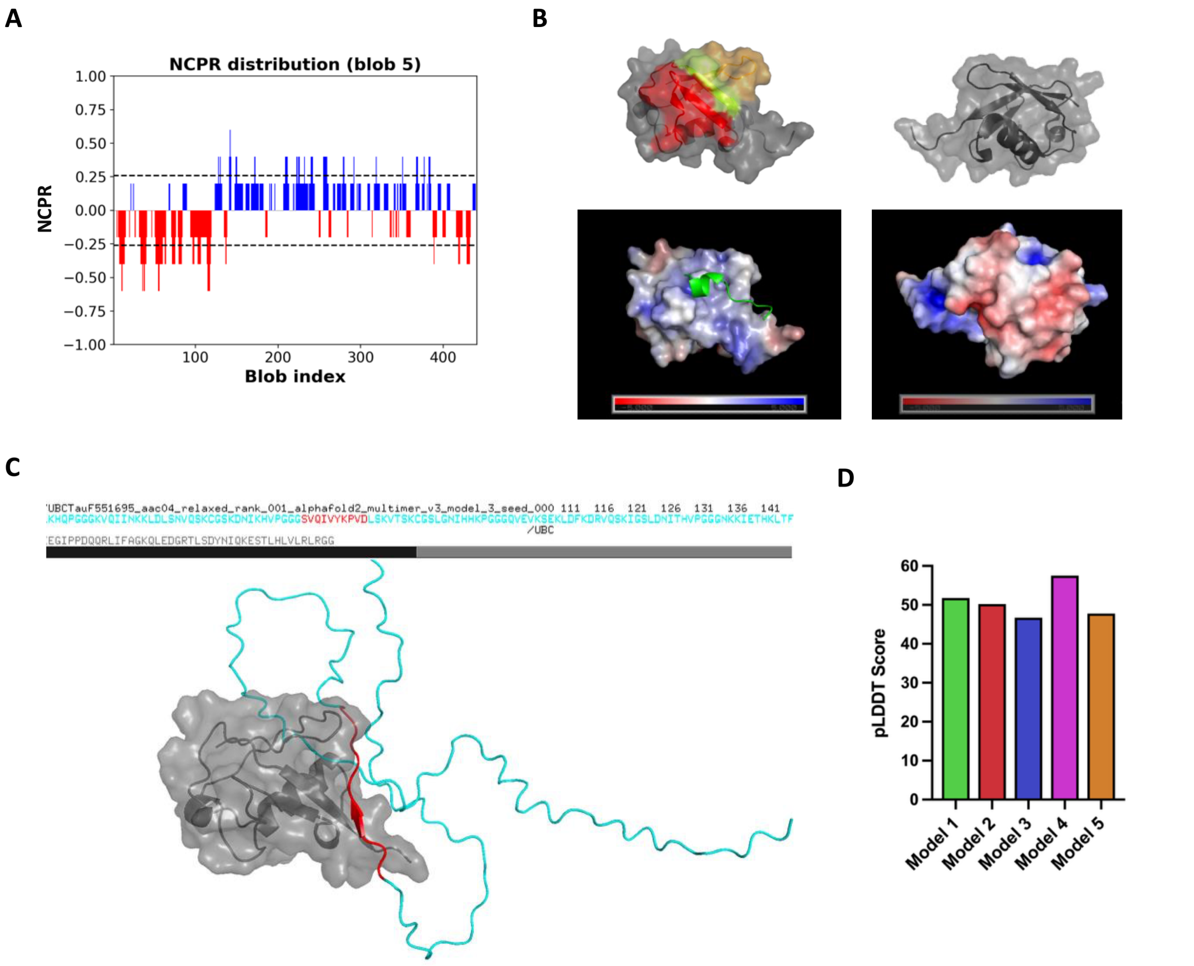


**Figure S4. Computational analyses predict Tau MTBR recognition of the ubiquitin hydrophobic patch.**

**(A–D)** Computational analyses independently support the microtubule-binding repeat (MTBR) region of Tau as the primary ubiquitin-binding interface. Charge distribution analysis of full-length Tau using CIDER identifies a positively charged central region encompassing the proline-rich domain and MTBR, flanked by negatively charged N- and C-terminal regions **(A)**. Mapping the disordered ubiquitin-binding motif (DisUBM) and residue charge distribution onto the ubiquitin structure highlights the canonical hydrophobic binding surface **(B)**. AlphaFold Multimer (v3) independently predicts an interaction between the Tau MTBR (cyan) and the ubiquitin (gray) hydrophobic patch. **(C)**. Confidence scores (predicted local distance difference test; pLDDT) for all five AlphaFold Multimer models are shown **(D)**.

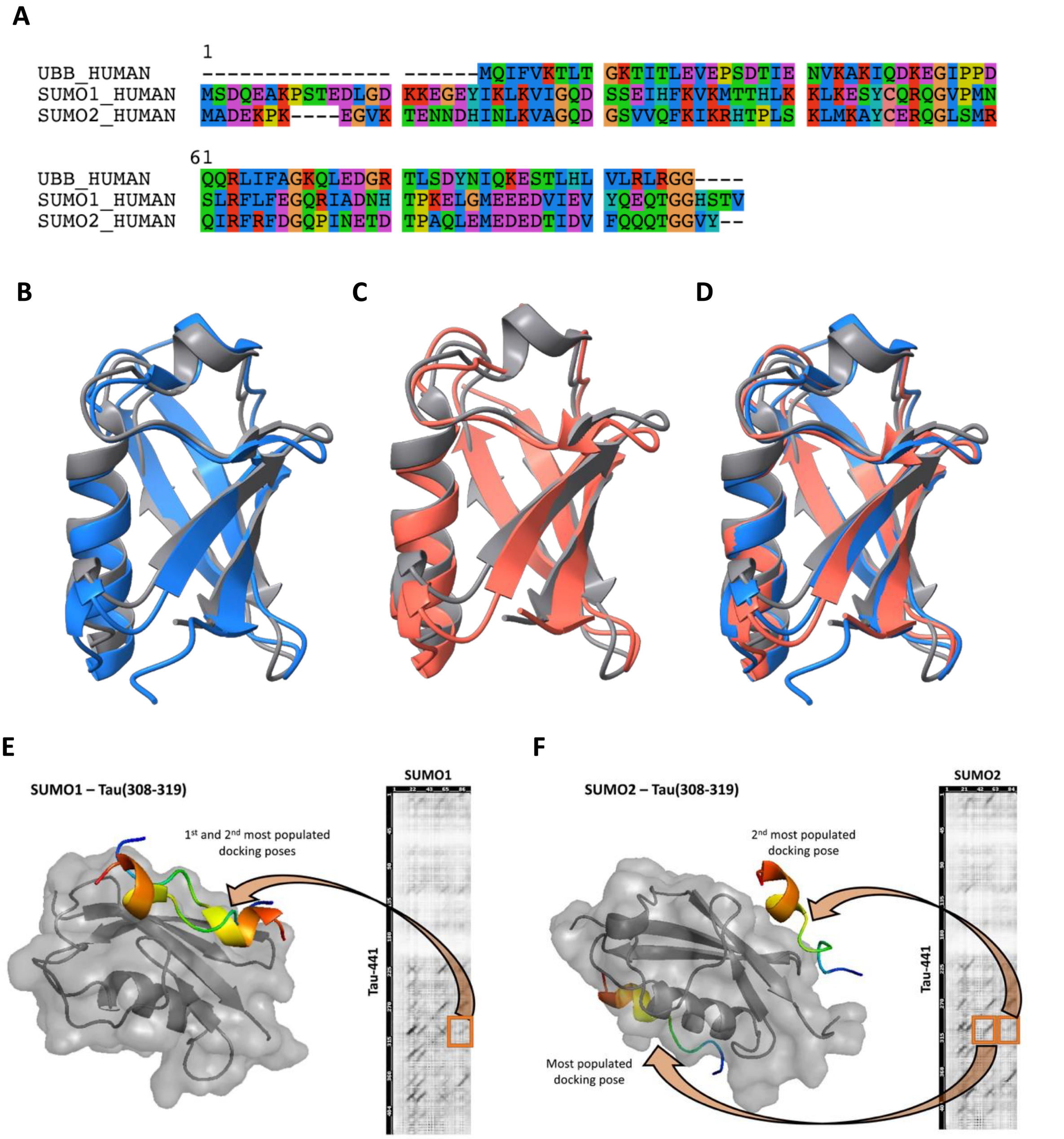


**Figure S5. Comparative structural analysis of ubiquitin and SUMO proteins supports a conserved mode of Tau recognition.**

**(A–F)** Comparative sequence and structural analyses suggest that the Tau ubiquitin-binding interface may extend to structurally related ubiquitin-like modifiers (UBLs). Sequence alignment and structural comparison of human ubiquitin (UBB), SUMO1, and SUMO2 reveal conserved and divergent structural features **(A–D)**. Structural overlays of SUMO1 (blue, PDB: 2UYZ) **(B)** and SUMO2 (salmon, PDB: 4BKG) **(C)** with ubiquitin (gray, PDB: 4XOF) **(D)**. Molecular docking and co-evolution analyses further predict that the Tau ubiquitin-binding region ($^{308}$IVYKPVDLSKVT$^{319}$; colored) can engage both SUMO1 **(**gray; **E)** and SUMO2 **(**gray; **F)** through interaction interfaces analogous to those observed for ubiquitin, providing structural support for a potentially conserved mechanism of Tau recognition among ubiquitin-like modifiers.

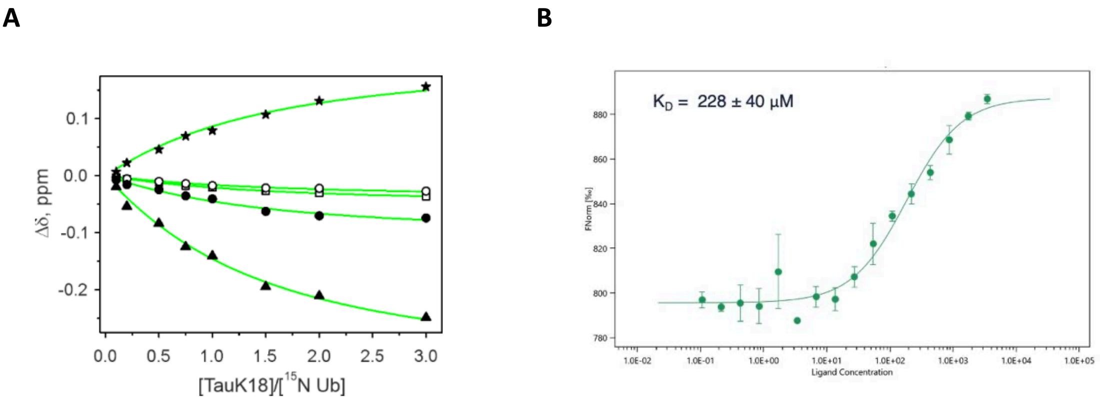


**Figure S6. Tau–Ub binding affinity determined by NMR and MST**

**(A)** Nuclear magnetic resonance (NMR) chemical shift titration of $^{15}$N-labeled ubiquitin (Ub) with Tau-MTBR (K18). Chemical shift perturbations (Δδ) of H (open symbols) and N (filled symbols) resonances of Ub backbone amides K6 (circles), A46 (squares), H68 (triangles), and L69 (stars) were fitted simultaneously to a 1:1 binding model. The solid lines show the best fit, yielding a shared $K_D$ value of 220 ± 39 µM. **(B)** Microscale thermophoresis (MST) independently validates the Tau-MTBR–ubiquitin interaction using 0.1 µM fluorescently labeled Tau-MTBR titrated with ubiquitin. Data represent three independent experiments performed at room temperature in 20 mM Tris-HCl pH 7.4, 150 mM NaCl, 10 mM $MgCl_2$, and 1 mM DTT.

movie 1: md1_0-440ns.mpg

movie 2: md2_0-540ns.mpg

**Movies M1 & M2. Molecular dynamics simulations support persistent multivalent interactions between Tau and diubiquitin.** Two independent molecular dynamics (MD) simulation replicas of the Tau–diubiquitin complex are shown in Figure 3. Tau-MTBR is shown in cyan, diubiquitin in gray, and interacting residues in orange/red (Tau) and yellow (diubiquitin). Replica 1 (MD1) dissociates after approximately 270 ns, whereas Replica 2 (MD2) remains associated longer, consistent with the interaction analyses shown in Figure 3E–F.

movie 3: syn_90ns.mpg

movie 4: syn_0-400ns.mpg

**Movies M3 & M4. Molecular dynamics simulations of the α-synuclein–SUMO1 complex.** Two independent MD simulation replicas of the α-synuclein (Syn)–SUMO1 complex reported by Panigrahi et al. (2023). Syn is shown in dark gray and SUMO1 in brown, with interacting residues highlighted in red (Syn) and yellow/green (SUMO1). These simulations illustrate a previously reported ubiquitin-like modifier interaction and are included for comparison with the Tau–Ubiquitin simulations shown in Supporting Movies M1 and M2.

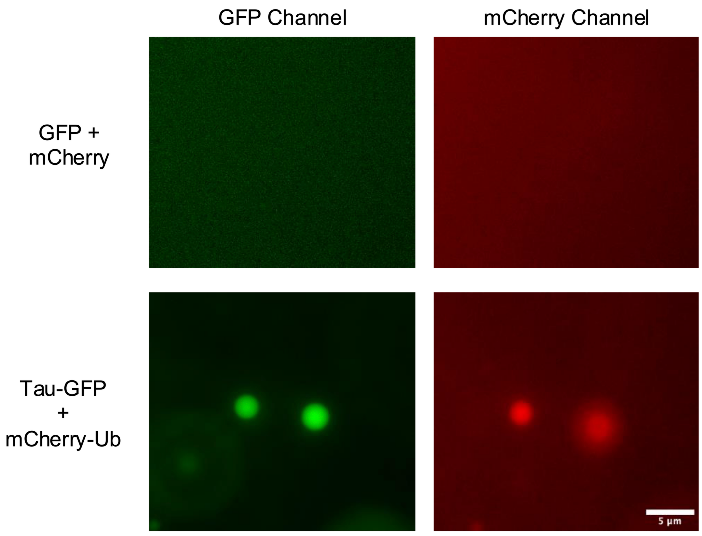


**Figure S7. mCherry-tagged ubiquitin partitions into Tau-GFP condensates.**
Representative fluorescence microscopy images showing enrichment of mCherry-ubiquitin within Tau-GFP condensates. GFP and mCherry alone remained diffuse under the same experimental conditions (10% PEG). GFP and mCherry fluorescence channels were acquired sequentially. Because Tau condensates move dynamically during image acquisition, the positions of individual droplets differ slightly between channels. Scale bar, 5 μm.

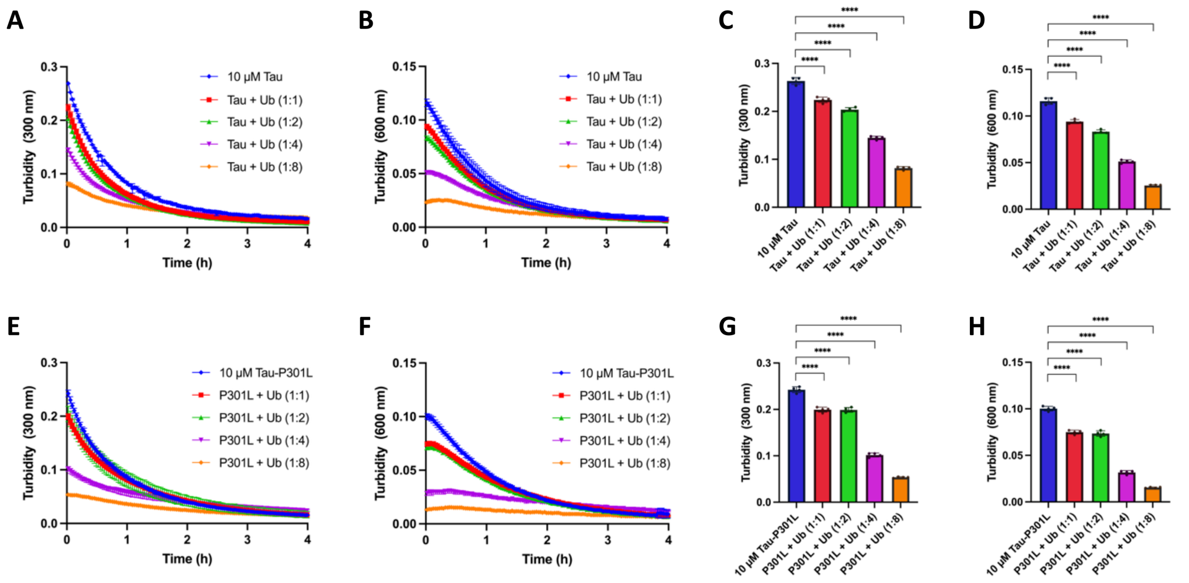


**Figure S8. Ubiquitin modulates Tau phase separation in a quantitative microplate assay.**
**(A–H)** The effects of mono-ubiquitin (Ub) on Tau phase separation were quantified by turbidity measurements. The turbidity of wild-type Tau **(A–D)** and Tau-P301L **(E–H)** was measured at 300 nm **(A, E)** and 600 nm (**B, F)** over 4 h at 25 ℃, in the absence or presence of increasing concentrations of monoubiquitin. Quantification of the maximum OD recorded throughout the assay at 300 nm **(C, G)** and 600 nm **(D, H)** demonstrates concentration-dependent suppression of phase separation for both Tau variants. Bar graphs represent mean ± SD (n = 4 technical replicates; one-way ANOVA with Holm–Sidak's multiple comparisons test; *****p* < 0.0001).

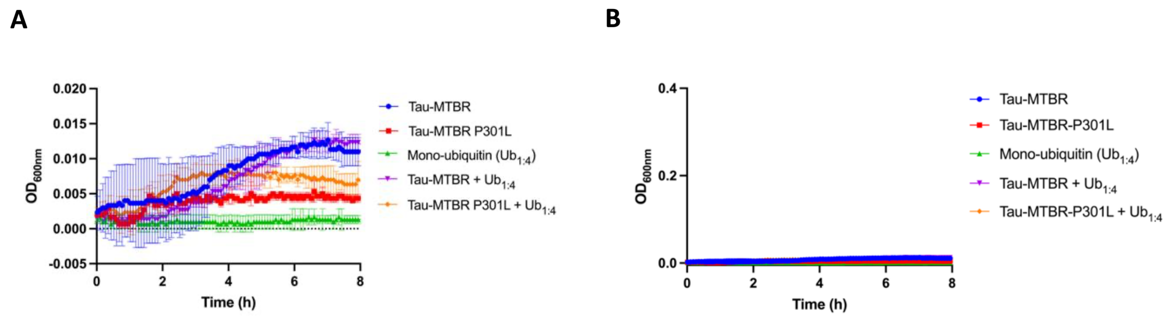


**Figure S9. Tau MTBR aggregation occurs in the absence of detectable liquid-liquid phase separation.**

**(A–B)** Optical density measurements at 600 nm recorded throughout the aggregation assay remained at baseline levels, indicating no detectable condensate formation under these experimental conditions **(A)**. The same data, displayed on the scale used for the LLPS turbidity assays **(B)** further demonstrate that the observed changes in Tau aggregation occurred independently of detectable LLPS. Graphs represent mean ± SD from three technical replicates.
